# SAIL: Sparse Autoencoders for Interpretable Alignment of Human Vision and Multimodal Large Language Models

**DOI:** 10.64898/2026.09.30.755515

**Authors:** Tan Gao, Luyuan Zhang, Yifei Wang, Yunjia Wang, Haotian Zhou, Mufan Xue, Guoyuan Yang

## Abstract

Alignment between human high-level visual representations and those of multimodal large language models (MLLMs) offers a quantitative framework for understanding information processing in human vision. However, similarities between model and brain representations do not directly reveal the semantic content underlying their correspondence, as this content is difficult to isolate in entangled MLLM representations. Here, we introduce SAIL, an interpretable framework that formulates voxel interpretation as a multi-unit sparse semantic correspondence problem between MLLMs and human vision. First, we train unified cross-layer sparse autoencoders (SAEs) on MLLM representations of Natural Scenes Dataset images and captions, and align each brain voxel with multiple SAE units. Second, we quantify semantic similarity between their top-response image sets to map semantic alignment across the cortex. We project multiple semantic labels back onto the cortex to interpret the content of this alignment. Finally, we assess these correspondences using category responses measured in independent fLoc experiments. Compared with SAEs trained independently at each layer, SAIL exhibits lower redundancy in SAE unit response profiles and higher semantic alignment. We find that semantic alignment is organized along the human ventral visual pathway across the models and subjects examined. The projected labels reveal semantic preferences within canonical visual regions, while fLoc experiments show agreement between predicted and measured category response patterns. Exploratory analyses further reveal cortical distributions of action-related semantics consistent with established functional accounts of lateral occipitotemporal and posterior parietal cortex. Overall, SAIL provides an interpretable account of the semantic correspondences between MLLM representations and human visual responses.

## 1 Introduction

Artificial neural networks have achieved remarkable success in modeling human brain responses, especially in high-level visual cortex (Wang et al., 2023; Chen et al., 2026; Schrimpf et al., 2021; Yamins et al., 2014; Gao et al., 2026). Building on this progress, multimodal large language models (MLLMs), with their integration of visual information and language context, offer a framework for probing high-level human visual processing (Jin et al., 2025; Wu et al., 2023; Caffagni et al., 2024). A central question is how MLLM representations correspond to the human visual system. Quantifying the semantic alignment between MLLMs and the human ventral visual pathway is one step in addressing this issue. Existing approaches relate MLLM predictions and representations to human similarity judgments and neural activity patterns, suggesting correspondences in object concepts and abstract emotion representations (Du et al., 2025; He et al., 2026). Other studies employ Representational Similarity Analysis (RSA) and encoding models to compare model representations with fMRI responses, revealing correspondences in representational structure and response patterns (Kriegeskorte & Kievit, 2013; Nili et al., 2014; Kriegeskorte et al., 2008a;b; Doerig et al., 2025; Ryskina et al., 2025). These studies establish model–brain correspondence, but leave its semantic content insufficiently understood.

Interpreting these correspondences requires addressing the entanglement of semantic information in model representations. Feature superposition encodes distinct semantic features along overlapping directions in representation space, so individual model units may exhibit polysemanticity, responding to multiple distinct concepts (Elhage et al., 2022; Olah et al., 2020). Matching such a unit to a voxel establishes a correspondence between their response patterns, but does not distinguish which of the unit’s semantic preferences accounts for that correspondence. Recent studies have explored residual decomposition (He et al., 2025) and sparse concept encoding (Zeng & Gallant, 2025) to make the semantic structure of brain-predictive representations more interpretable.

Sparse autoencoders (SAEs) have emerged as a data-driven tool for separating interpretable response patterns from entangled model activations (Huben et al., 2024; Templeton, 2024). Like other sparse coding methods, they learn overcomplete dictionaries (Lee et al., 2006); empirical studies have examined their interpretability in language and vision-language models (Rajamanoharan et al., 2024; Pach et al., 2025), while theoretical work has established feature recovery guarantees under specific assumptions (Chen et al., 2025). SAEs have also been used to align deep visual representations with fMRI responses and reveal hierarchical correspondences along the human ventral visual pathway Mao et al. (2026). Yet interpretable units alone do not resolve how to characterize multiple semantic preferences within a voxel.

Existing frameworks have explored multi-semantic descriptions of visual cortex (Yang et al., 2025; Gao et al., 2026). SAE-BrainMap assigns each voxel a single best-matching SAE unit, providing a semantic description based on that unit Mao et al. (2026) This characterizes a dominant preference but does not describe the range of semantic content associated with other matched units. A semantic account of such responses therefore motivates the examination of multiple matched units rather than restricting the explanation to one. However, simply increasing the number of matches may not provide more distinct semantic content. When SAEs are trained independently at each layer, units of different layers can respond to similar images and express overlapping meanings. A voxel matched to several such units may therefore receive repeated descriptions of similar response preferences. Therefore, when matching a voxel to multiple semantic units, we also need to consider the degree of redundancy among their response patterns. Previous work has integrated representations from multiple layers for visual tasks and neural encoding (Wen et al., 2018; Hariharan et al., 2015). Building on this approach to combining hierarchical information, we jointly learn sparse units across layers and evaluate their activation redundancy and semantic alignment with voxel response preferences.

In this work, we address these challenges in interpreting brain–model alignment by introducing SAIL, an interpretable framework that formulates voxel interpretation as a multi-unit sparse semantic correspondence problem between MLLMs and human vision, yielding per-voxel interpretable semantic units. We integrate representations from multiple MLLM layers and train a unified cross-layer SAE, replacing independent decomposition at each layer with joint learning in a shared feature space. Each voxel is then matched to multiple units, and semantic alignment is evaluated by comparing the image sets that elicit their highest responses. We utilize the Natural Scenes Dataset (NSD) (Allen et al., 2022) as a primary benchmark, providing MLLMs with the natural images viewed by the subjects and their associated captions. The captions provide linguistic context for the images, allowing us to examine how the resulting multimodal representations align with measured human brain responses. We compare unit redundancy and semantic alignment against layer-wise SAEs, with single unit matching, vision tower, and projector controls examining alternative configurations. Searchlight RSA provides a complementary evaluation metric. Cortical label maps characterize semantic organization, independent fLoc responses assess category response reconstruction (Stigliani et al., 2015), and exploratory analyses examine action-related semantics beyond predefined selective ROIs.

## 2 Methods

### 2.1 Dataset

We used the NSD to study semantic alignment between human vision and MLLMs (Allen et al., 2022). NSD provides high-resolution fMRI responses from eight participants viewing natural images. Beta estimates were prepared with the *betas fithrf GLMdenoise RR* pipeline, z-scored across runs, and averaged across repeated presentations to obtain one response per voxel per image. Cortical surfaces were reconstructed using FreeSurfer (Fischl et al., 1999; Dale et al., 1999), and stimuli consisted of square-cropped COCO images subtending 8.4° × 8.4° of visual angle (Lin et al., 2014). Related preprocessing choices have been adopted in prior NSD studies (Xue et al., 2024; Yang et al., 2025).

### 2.2 Region of Interest

In addition to whole-brain semantic alignment, we focused on a set of ventral visual ROIs. These included early visual areas, as well as higher-level category-selective visual regions, including the fusiform face area (FFA) for face selectivity (Kanwisher et al., 1997), extrastriate body area (EBA) for body selectivity (Downing et al., 2001), retrosplenial cortex (RSC) for place selectivity (Brodmann, 1909), visual word form area (VWFA) for word selectivity (Cohen et al., 2000), and the food-selective region (FOOD) (Jain et al., 2023).

### 2.3 SAIL FRAMEWORK

SAIL recasts voxel-level alignment as an interpretable semantic correspondence between cortical voxels and sparse semantic units within the model. The framework first learns sparse units from multiple MLLM layers, then matches these units to cortical voxels, and finally evaluates the semantic agreement of the resulting correspondences on held-out images (Figure 1). We train and evaluate SAIL across eight subjects using four MLLMs: LLaVA-v1.6-Mistral-7B, LLaVA-v1.6-Vicuna-7B, DeepSeek-VL-7B-Chat, and Qwen2.5-VL-7B-Instruct (Liu et al., 2023; 2024; Lu et al., 2024; Bai et al., 2025). Unless otherwise noted, main-text results are presented for Qwen2.5-VL-7B-Instruct and subject S5; results for the remaining models and subjects are reported in the Appendix.

**Figure 1:**
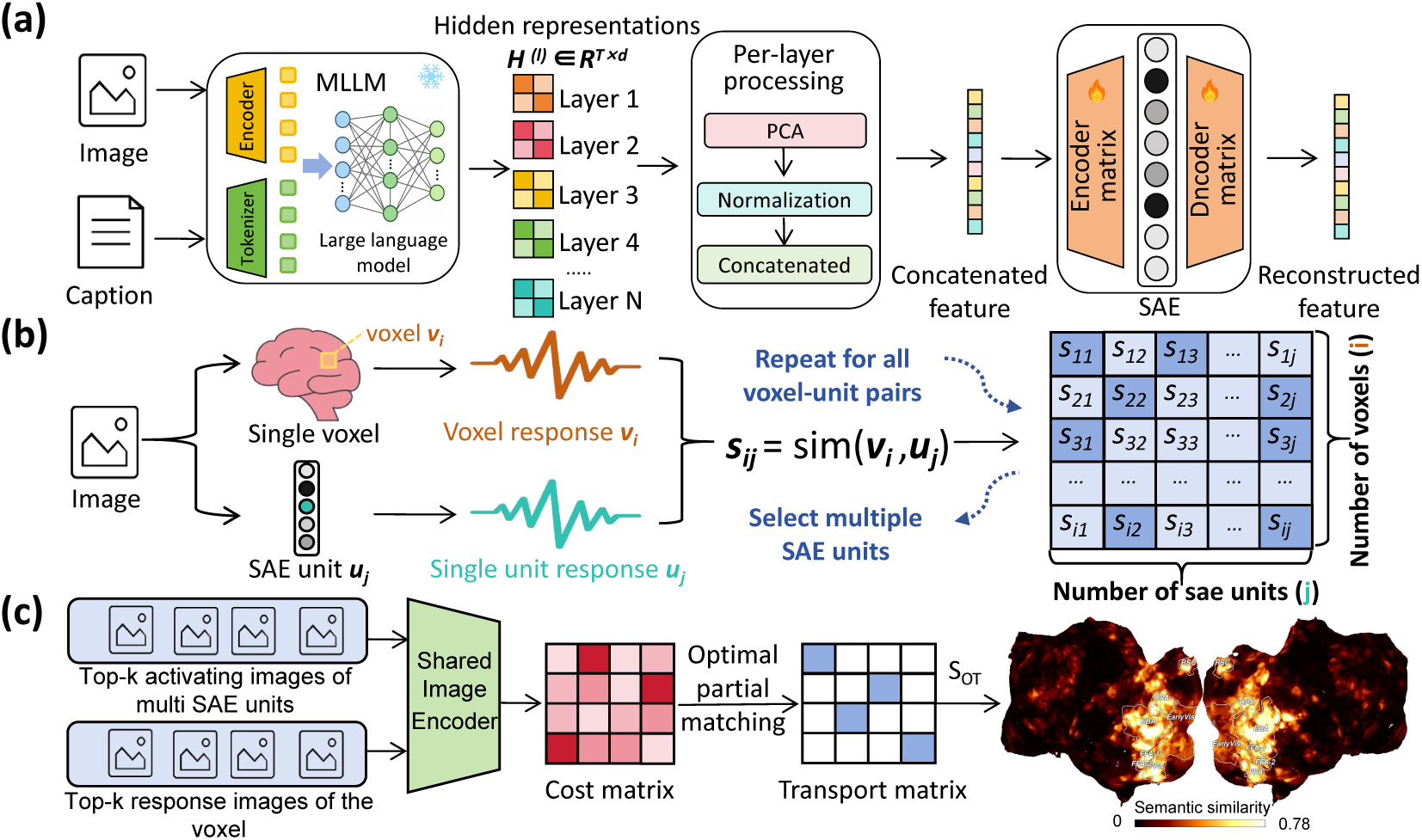
Architecture of SAIL. **(a)** Each image–caption pair is processed by an MLLM. Layerwise features are transformed and concatenated to train a unified cross-layer SAE. **(b)** For each voxel, response patterns are matched to SAE unit activation patterns, yielding multiple matched units. **(c)** SEED-based image similarities and Optimal Transport (OT) quantify semantic agreement between voxel preferred images and images preferred by the matched SAE units. The resulting scores are mapped onto the cortical surface.

#### Training Unified Cross-Layer SAE

Given a natural image and its caption, we extract representations from multiple MLLM layers, apply PCA within each layer (Wen et al., 2018), and standardize the components to comparable scales (Laturnus et al., 2020). We concatenate these representations for unified SAE training. In five-fold cross-validation, preprocessing and SAE training use only the training split, with semantic alignment evaluated on held-out images.

Given a concatenated representation **x**, the SAE encodes it into a nonnegative sparse activation a and reconstructs the input:

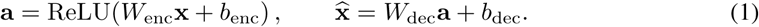

Here, x, a, and x denote the concatenated input, sparse activation, and reconstruction, respectively; *W* and *b* are learned parameters. Decoder columns are normalized to unit norm during training. To retain information from the input while encouraging sparse activations, we minimize

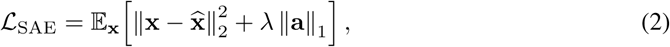

where *λ* controls the trade-off between reconstruction fidelity and activation sparsity (Cunningham et al., 2023; Bricken et al., 2023). Further details are provided in Appendix A .

#### Matching Voxel to SAE Units

Having learned the sparse units, we next identify those whose activation patterns correspond to each voxel’s response pattern. Matching uses the training images within each fold. Let **v***_i_* and **u***_j_* denote the response vector of voxel *i* and the activation vector of SAE unit *j* across these images. After *L*_2_ normalization, we use cosine similarity as the matching score:

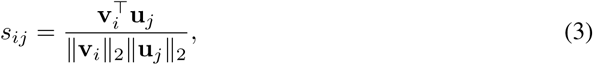

Repeating this computation for all voxel–unit pairs yields a similarity matrix **S**. Rather than assuming a one-to-one mapping between voxels and units, we rank SAE units by *s_ij_* for each voxel and retain the *K* highest-scoring units. These units provide a joint interpretation of the voxel’s semantic preferences.

#### Quantifying Semantic Alignment

To assess whether response-pattern matches reflect semantic agreement, we compare two image sets from each fold’s held-out pool: the voxel’s *K* highest-response images, *I_vi_* = *{p*_1_*, . . ., p_K_}*, and the highest-activating image for each of its *K* matched units, *I_ui_* = *{q*_1_*, . . ., q_K_}*. The SAE-preferred images are represented as a collection of *K* image entries. We evaluate set-level agreement using pairwise semantic similarities rather than individual images or averaged embeddings. For the pairwise comparison, we use SEED semantic similarity (Park et al., 2026) to quantify semantic similarity between individual images from the two preferred image sets. Here, *p_a_ ∈ I_vi_* denotes the *a*-th image in the voxel preferred image set, and *q_b_ ∈ I_ui_* denotes the *b*-th image in the SAE preferred image set. The metric combines object-level agreement, caption similarity, and visual feature similarity. *S*_obj_ measures agreement between automatically detected object categories using the SEED Object F1 score. *S*_cap_ computes cosine similarity between embeddings of captions generated from the two images. *S*_eff_ computes the correlation between image feature vectors extracted by a pretrained EfficientNet (Tan & Le, 2019). Following the original SEED formulation, we average the three components to obtain the semantic similarity of each image pair:

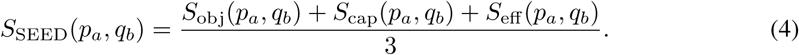

The evaluation models and implementation details are provided in Appendix B. For each voxel, pairwise SEED similarities define a *K × K* semantic cost matrix:

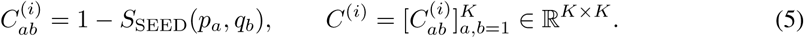

A smaller 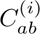 indicates greater semantic agreement between the two images. To obtain a set-level comparison, we next determine which images correspond and aggregate their matching costs. One approach is to pair images by their positions, but this makes the comparison depend on an arbitrary ordering. Another approach is to select the lowest-cost match independently for each voxel-preferred image, but this may repeatedly select the same SAE-preferred image. We therefore use discrete optimal transport (OT) to determine the correspondences jointly. We solve for the optimal transport plan *P*\* that minimizes the total semantic cost (Chapel et al., 2020; Cuturi & Peyré, 2019):

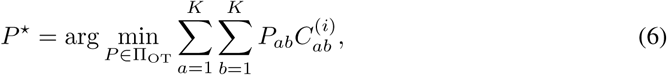

where Π_ΟΤ_ is the feasible set of transport plans (constraints in Appendix C), and *P_ab_* is the transport mass assigned to (*p_a_, q_b_*). We convert the optimal transport cost into voxel–unit semantic similarity:

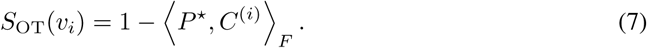

Higher scores indicate greater semantic agreement between the two image sets. We average scores across the five held-out folds to obtain each voxel’s final similarity.

## 3 Redundancy in SAE Unit Response Profiles

When SAEs are trained independently at each layer, reconstruction and sparsity are optimized separately, without a joint objective coordinating features across layers. Similar response profiles may therefore recur in different layer-specific dictionaries, causing multiple matched units to provide repeated descriptions of a voxel. SAIL instead learns a unified dictionary from concatenated layer representations under a common reconstruction objective and sparsity penalty. We therefore first compare redundancy in SAE unit response profiles between SAIL and SAEs-BrainMap to examine whether this joint formulation reduces repeated response patterns. Figure 2a schematically illustrates the difference in unit redundancy between independently trained layer-specific SAEs and a unified cross-layer SAE. To assess unit-level representational redundancy, we represented each active unit by its activation profile over a common set of held-out images. Its redundancy score was defined as the maximum cosine similarity to any other candidate unit within the same subject and method. Candidate units were pooled across layers for SAEs-BrainMap and drawn from the unified SAE for SAIL. Across all eight subjects, SAEs-BrainMap showed higher redundancy distributions than SAIL (Figure 2b), indicating greater duplication of unit response patterns. To control for differences in candidate unit counts, we performed 200 matched-size unit resamples and compared the mean redundancy between methods in each resample. The resulting SAEs-BrainMap-minus-SAIL difference distributions were shifted toward positive values for all eight subjects (Figure 2c), supporting the stability of SAIL’s lower redundancy under the tested resampling procedure. Results for the remaining three MLLMs are provided in Appendix D.

**Figure 2:**
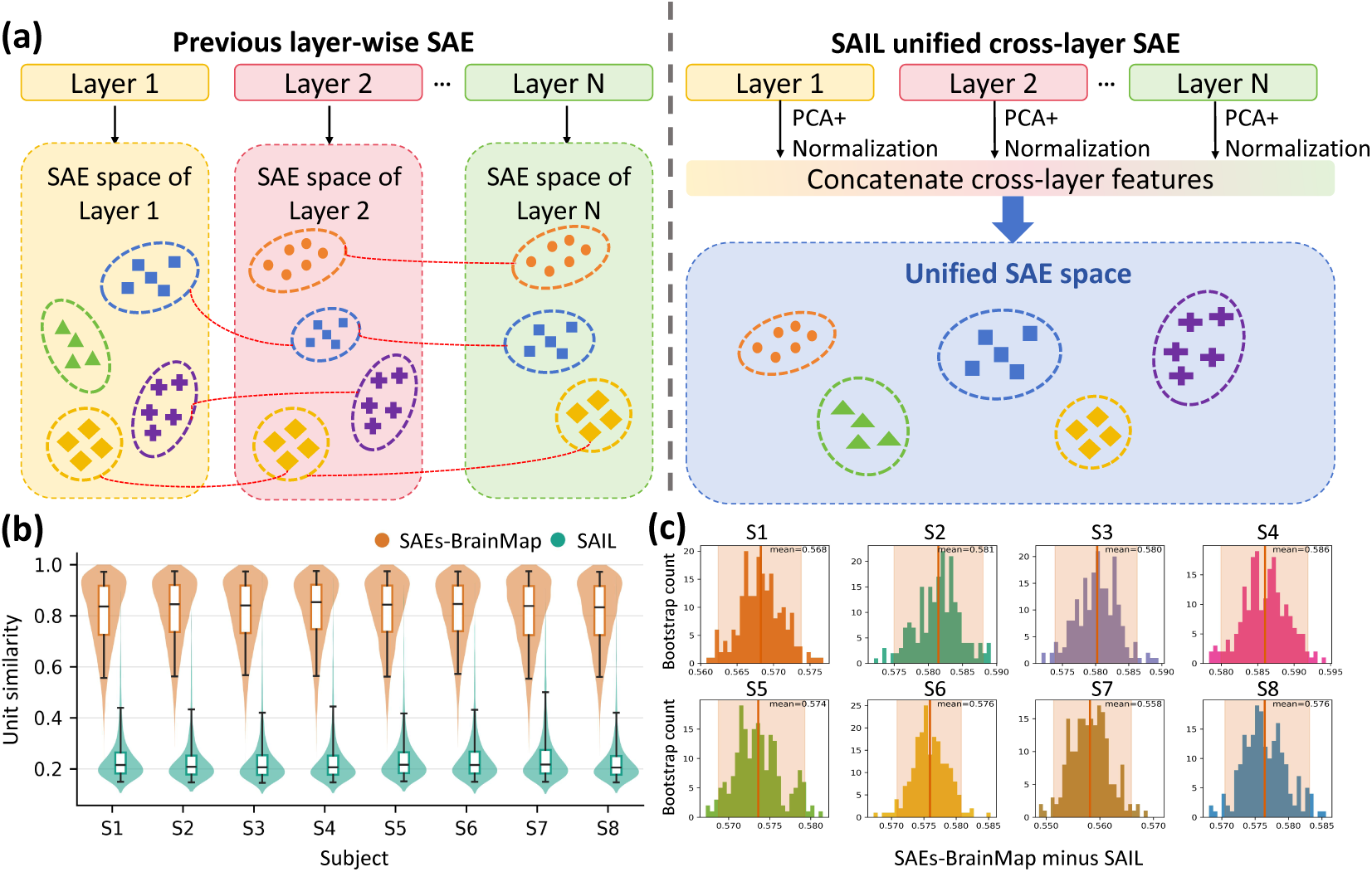
Unified cross-layer SAE reduces response redundancy across units. **(a)** Previous layer-wise SAEs learn independently at each layer and can produce similar response profiles in units from different layers. SAIL trains a unified SAE on concatenated cross-layer features, allowing units to be learned in a shared space. **(b)** Distributions of maximum cosine similarity between SAE unit response profiles for SAEs-BrainMap and SAIL across eight subjects. Both methods use matched numbers of active units and are evaluated on the same held-out images. **(c)** Distributions of the SAEs-BrainMap-minus-SAIL mean maximum unit similarity from 200 matched-size resamples. Lines denote means and shading denotes 95% intervals.

## 4 Semantic Alignment Comparison

The preceding analysis shows that unified cross-layer training reduces redundancy in SAE unit response profiles. We next examine whether this less repetitive response organization is accompanied by higher semantic alignment between the model and the brain. Figure 3a maps the cortical distribution of SAIL semantic similarity, while Figure 3b compares methods across eight subjects over the whole cortex and within selective ROIs. SAIL achieves higher semantic similarity averaged across subjects in both evaluation masks than SAEs-BrainMap, which trains SAEs independently at each layer. SAIL thus exhibits not only lower redundancy in SAE unit response profiles but also closer semantic correspondence with voxel response preferences.

**Figure 3:**
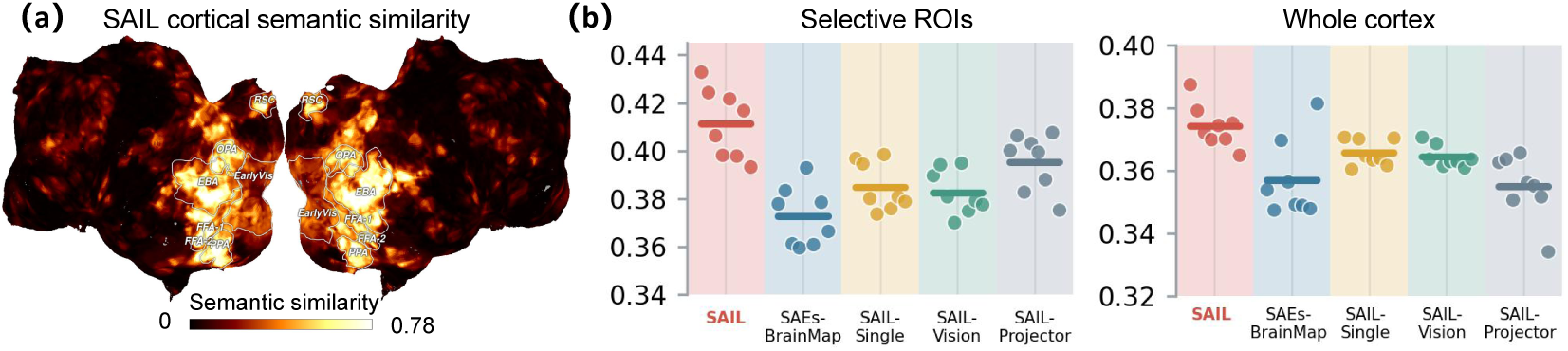
Cortical semantic alignment with SAIL. **(a)** Cortical distribution of SAIL semantic similarity for Subject S5. **(b)** Semantic similarity in the Whole cortex and Selective ROI of eight subjects, comparing SAIL, SAEs-BrainMap, SAIL-Single, SAIL-Vision, and SAIL-Projector. Each point represents the mean semantic similarity for one subject; horizontal lines indicate the mean across eight subjects.

We next compare semantic alignment when matching a single unit per voxel or using visual representations alone (Figure 3b). SAIL-Single uses the same language-layer representations, held-out image pool, and semantic evaluation procedure as SAIL, but restricts each voxel to its best-matching unit. Its lower semantic alignment indicates that matching multiple units provides a more informative description of voxel response preferences than single-unit matching. SAIL-Vision and SAIL-Projector use outputs from the vision tower and projector, respectively, whereas the full SAIL representation is derived from the multimodal input containing the image and its caption. Both controls yield lower semantic alignment than SAIL, supporting the use of multimodal representations for characterizing voxel response preferences.

Results for the remaining three MLLMs are provided in Appendix E. We further compare SAIL with SAEs-BrainMap using searchlight RSA under the same matching procedure. This provides an additional comparison based on representational structure rather than the semantic similarity of preferred image sets. See Appendix F for the evaluation settings and additional results.

## 5 Interpreting Cortical Semantic Alignment

The preceding analyses assessed the degree of semantic alignment between model representations and brain responses. We next examine the semantic content underlying these correspondences and its spatial organization across the cortex. For each matched SAE unit, we extract candidate labels from the captions of its maximally activating image. Keeping the image fixed, we pair it with each candidate label in a common text template and compute the activation of the target SAE unit through the same MLLM–SAE pipeline. The candidate label eliciting the highest activation is selected as the semantic label. Each voxel thus receives multiple labels from its matched units. Details are provided in Appendix H. To examine the cortical organization of these labels, we visualize each voxel’s multi-label description in a continuous image-semantic space. Drawing on the UMAP visualization procedure of BrainSCUBA (Luo et al., 2024), we extend it to a similarity-weighted multi-label representation. We first compute CLIP image embeddings for all NSD stimulus images and fit a three-dimensional UMAP to these representations. We then encode the labels using the corresponding CLIP text encoder and project their embeddings into the fitted UMAP space.

For each voxel, we average the UMAP coordinates of its multiple semantic labels, weighted by their normalized semantic alignment scores, and convert the resulting coordinates to RGB. Figure 4a visualizes the spatial distribution of these multi-label representations in higher-order visual cortex. Figure 4b provides a lexical interpretation of the UMAP directions: one direction is dominated by person-related words, including *person*, *people*, and *player*; a second contains scene or spatial words, including *kitchen*, *room*, and *street*; and a third contains food-related words, including *food*, *cake*, and *sausage*. The map places these semantic representations in relation to established category-selective regions. Person- and body-related semantics are displayed near EBA and FFA, scene-related semantics near RSC, OPA, and PPA, and food-related semantics in the food-selective territory adjacent to FFA. Complementing the cortical maps, Figure 4c presents semantic interpretations of individual voxels. Each label corresponds to a matched SAE unit and is shown with its highest-activating image. See Appendix G for additional cortical semantic maps.

**Figure 4:**
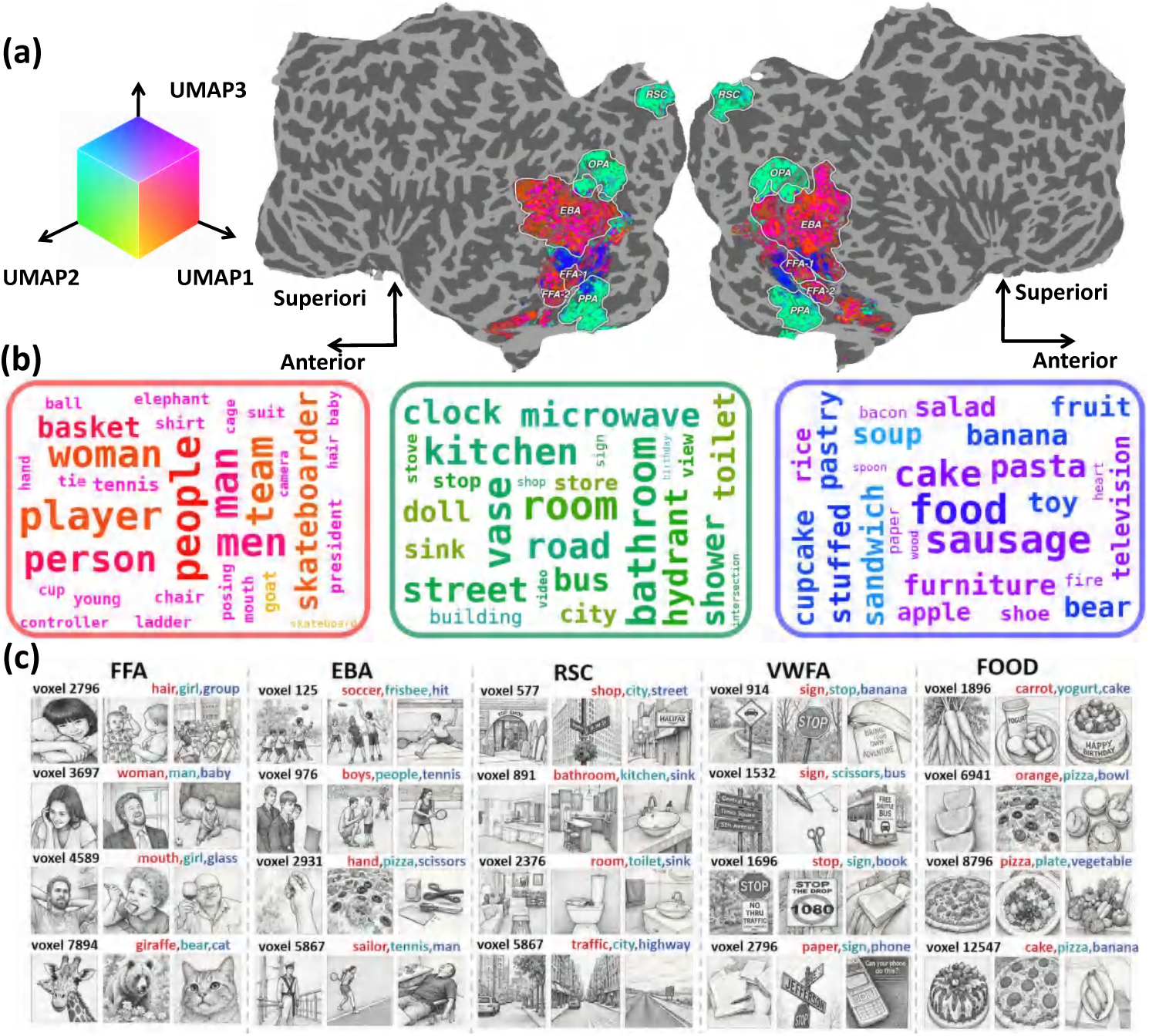
UMAP visualization of multi-label semantics on cortex. **(a)** A three-dimensional UMAP is fit to CLIP image embeddings from all NSD stimulus images. SAIL labels are embedded using the CLIP text encoder and projected into the fitted UMAP space. For each voxel, the coordinates of multiple semantic labels are combined using normalized semantic alignment scores, converted to RGB values, and projected onto higher-order visual cortex. **(b)** Representative words along the three UMAP directions. The word clouds include only labels observed in the SAIL results; word color is determined by UMAP coordinates, and word size indicates directional similarity to the corresponding UMAP axis. **(c)** Semantic interpretations for selected voxels in FFA, EBA, RSC, VWFA, and FOOD. Each example includes a voxel identifier, multiple semantic labels, and images that strongly activate the corresponding matched SAE units.

## 6 fLoc Category Response Reconstruction

Building on the regional organization of SAIL semantic maps, we test whether SAIL reconstructs category response patterns in an independent fLoc experiment (Stigliani et al., 2015). Keeping voxel–unit correspondences fixed, we average unit activations within each category and predict each voxel response by weighting its matched unit activations with normalized matching scores. We first compare responses to bodies and places. The SAIL predicted body–place contrast and the measured fLoc contrast show similar regional organization (Figure 5a). Both maps show body preference near EBA and place preference in scene-selective regions, including RSC, OPA, and PPA. Figure 5b shows representative stimuli used in this comparison. These results show that the voxel–unit correspondences established by SAIL enable the reconstruction of regional response differences between body and scene categories in an independent fLoc experiment. We next move from differences between regions to response patterns within a region. Figure 5c compares SAIL predicted and measured responses to limb images in bilateral EBA for subject S5. The maps show similar broad spatial structure, with several high-response locations appearing in corresponding parts of EBA. Each map was normalized separately within EBA for visualization. Table 1 summarizes category response reconstruction across eight fLoc categories and eight subjects, showing correspondence between SAIL predictions and measured responses. These results extend the evaluation from cortical semantic organization to stimulus-evoked response patterns, indicating that SAIL-derived voxel–unit correspondences remain informative in an independent fLoc experiment. Results for the remaining three MLLMs are provided in Appendix I.

**Figure 5:**
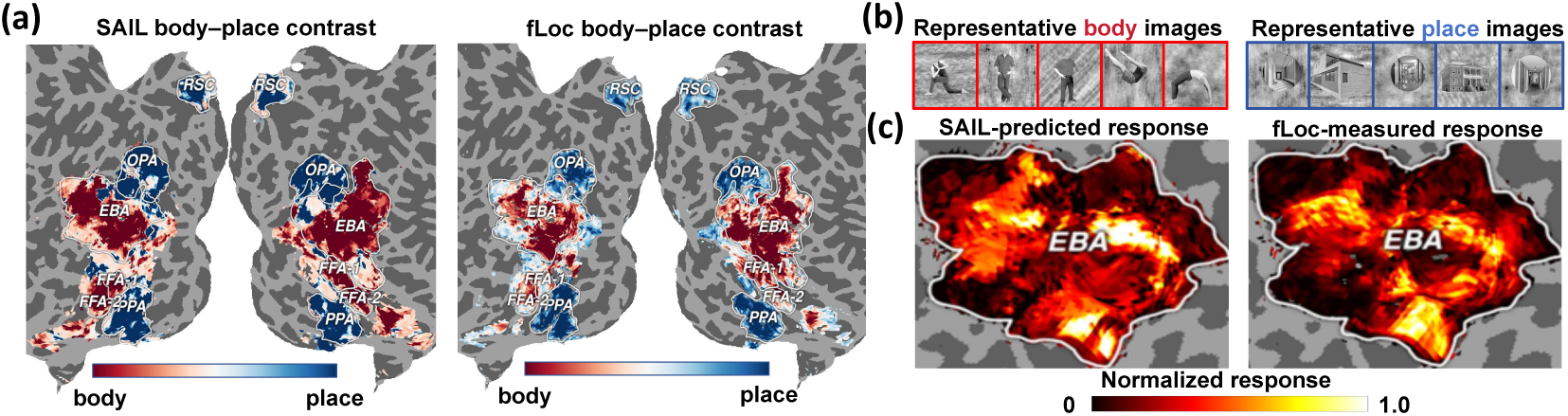
Correspondence between SAIL predictions and measured fLoc category responses. **(a)** SAIL predicted and fLoc measured body–place contrasts for subject S5. Warm colors indicate body preference, and cool colors indicate place preference. **(b)** Representative stimuli from the fLoc body and place categories. **(c)** SAIL predicted and fLoc measured limb responses in bilateral EBA. The two maps are normalized separately within EBA to compare their spatial distributions.

**Table 1:** Cosine similarity between SAIL predicted and measured fLoc category responses.

| Subject | fLoc category |  |  |  |  |  |  |  |
| --- | --- | --- | --- | --- | --- | --- | --- | --- |
|  | Adult | Body | Child | Corridor | House | Limb | Number | Word |
| S1 | 0.784 | 0.683 | 0.759 | 0.691 | 0.725 | 0.739 | 0.731 | 0.828 |
| S2 | 0.789 | 0.722 | 0.833 | 0.761 | 0.713 | 0.747 | 0.661 | 0.796 |
| S3 | 0.810 | 0.717 | 0.847 | 0.778 | 0.696 | 0.790 | 0.751 | 0.834 |
| S4 | 0.826 | 0.728 | 0.789 | 0.742 | 0.647 | 0.723 | 0.533 | 0.759 |
| S5 | 0.802 | 0.671 | 0.794 | 0.753 | 0.704 | 0.742 | 0.647 | 0.815 |
| S6 | 0.797 | 0.693 | 0.813 | 0.781 | 0.685 | 0.709 | 0.656 | 0.800 |
| S7 | 0.762 | 0.678 | 0.813 | 0.738 | 0.621 | 0.762 | 0.651 | 0.805 |
| S8 | 0.813 | 0.768 | 0.848 | 0.817 | 0.745 | 0.750 | 0.653 | 0.812 |
| **Mean** | **0.798** | **0.708** | **0.812** | **0.758** | **0.692** | **0.745** | **0.660** | **0.806** |

## 7 Investigating the Brain’s Action Representations

Having validated SAIL within selective ROIs, we further explore whether it can reveal meaningful cortical semantic organization beyond these predefined regions. We focus on action-related semantics, which connect body posture, object interactions, and action goals in natural images. In addition to body-selective regions, action information is represented across a distributed system encompassing lateral occipitotemporal and posterior parietal cortex (Caspers et al., 2010; Lingnau & Downing, 2015; Tucciarelli et al., 2019). We ask whether SAIL’s multi-semantic labels can characterize this broader cortical organization using static natural images.

We identified action-related labels using a predefined vocabulary of 106 terms spanning movement, posture, sports, and interactions between people and objects. For each voxel, we quantify the weighted proportion of action-related labels among its multiple semantic labels, using semantic alignment scores as weights to measure the relative contribution of action semantics. Figure 6a displays the cortical map for subject S5, while Figure 6b provides ventral and bilateral lateral views. The vocabulary and its construction are provided in Appendix K. The underlying action-semantic scores show elevated values in bilateral lateral occipitotemporal cortex (LOTC) and parts of posterior parietal cortex, extending towards superior parietal lobule (SPL) and inferior parietal lobule (IPL). This distribution is consistent with previous accounts of action representation. LOTC represents information about body and tool movements and action meaning, with action representations organized along dimensions including sociality and transitivity (Lingnau & Downing, 2015; Wurm et al., 2017; Tucciarelli et al., 2019; Kabulska et al., 2024). Posterior parietal cortex also contributes to action processing (Caspers et al., 2010): SPL differentiates among classes of observed actions (Abdollahi et al., 2013), whereas IPL supports action representations that generalize across specific visual instances (Wurm & Lingnau, 2015). These correspondences extend semantic characterization beyond predefined selective ROIs. Importantly, this analysis provides an exploratory localization of action-related semantics rather than an independent demonstration of action selectivity.

**Figure 6:**
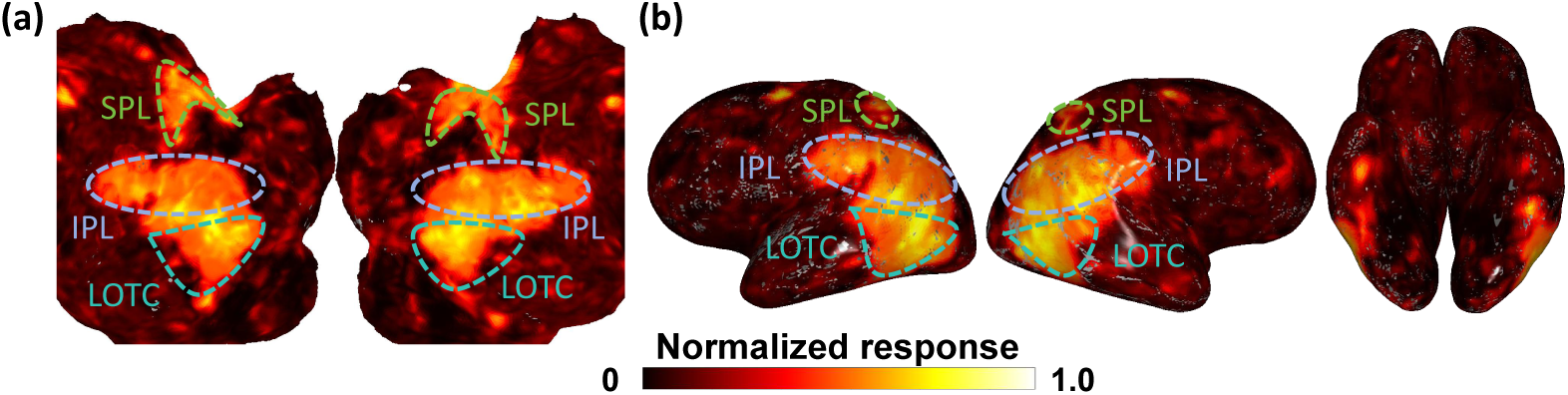
Cortical distribution of action-related multi-semantic labels. **(a)** Cortical distribution of SAIL action-semantic scores in subject S5, shown on a flattened surface with LOTC, SPL, and IPL outlined for anatomical reference. **(b)** Corresponding surface renderings from left lateral, right lateral, and ventral perspectives.

## 8 Discussion

The value of SAIL lies not only in stronger semantic alignment, but also in making the content of that alignment interpretable. Unified cross-layer training reduces redundancy relative to layerwise SAEs, while multiple matched units describe voxel preferences beyond a single dominant concept. Cortical maps and independent fLoc responses connect these interpretations to regional organization and category responses, offering an interpretable description of brain–model correspondence beyond overall similarity.

Several limitations remain. Preferred images and labels characterize semantic preferences within the evaluated stimulus set, but do not fully distinguish individual concepts from co-occurring visual content. Controlled stimulus comparisons could further distinguish the semantic preferences described by these labels from co-occurring visual content. Moreover, the present analysis focuses on static images; dynamic stimuli would allow examination of how brain–model correspondences evolve over time.

## A Preprocessing and SAE Training

### Data and preprocessing

We partition each subject’s stimulus records into five folds, using four folds for training and the remaining fold for held-out evaluation. Within each training split, records with non-finite fMRI responses are excluded from voxel–unit matching together with the corresponding unit activations. All models use the same subject-specific splits, generated with seed 42 + *s*, where *s* denotes the subject index. With MLLM parameters fixed, we average each language layer’s outputs across the multimodal input sequence. Within each fold, we fit randomized PCA separately to each selected layer using only the training images and retain 256 principal components per layer. We then standardize the retained components using statistics estimated from the training split and concatenate them in layer order. The fitted PCA transformations and standardization parameters are applied unchanged to the held-out images.

The selected language layers have zero-based indices 0–27 for Qwen2.5-VL-7B-Instruct, 0–29 for DeepSeek-VL-7B-Chat, and 0–31 for both LLaVA-v1.6-Mistral-7B and LLaVA-v1.6-Vicuna-7B. We use the publicly released checkpoints listed above and keep all MLLM parameters frozen throughout the analysis.

### SAE training

The SAE comprises the ReLU encoder and linear decoder defined in the main text, with a hidden dimension four times the concatenated input dimension. After each optimization step, each decoder column is renormalized to unit Euclidean norm. For every model, we use Adam with a learning rate of 5 *×* 10*^−^*^5^ and train for 5,000 updates per fold. Each batch contains 4,096 training examples sampled with replacement. The loss combines mean squared reconstruction error with 5.0 times the mean absolute activation. Because the hidden dimension is four times the input dimension, this objective is equivalent, up to an overall multiplicative constant, to the unnormalized norm formulation in the main text with *λ* = 1.25. We implement preprocessing and training with scikit-learn and PyTorch, respectively, and train each SAE in float32 on a single NVIDIA L20 GPU.

### Variance retained by PCA

We retain 256 principal components per layer to control the dimensionality of the concatenated representation. We assess variance retention using the explained variance ratios stored in the fitted PCA models. For each model, we summarize cumulative explained variance across all selected language layers, eight subjects, and five folds. The median variance retained by 256 components ranges from 87.49% to 93.84% across models (Table A.1), showing that this representation preserves most of the training feature variance. Figure A.1 traces the accumulation of explained variance over the retained components.

**Table A.1:**
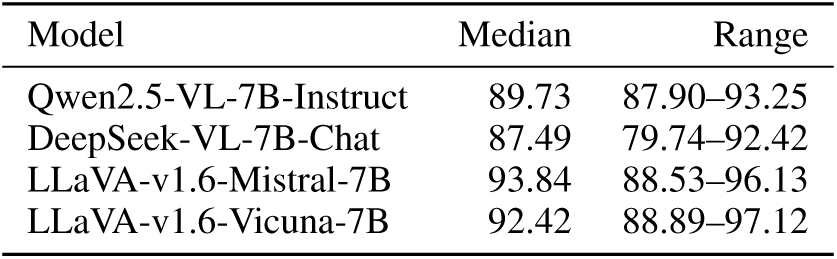
Cumulative explained variance of 256 principal components per layer (%). Median and range summarize all selected language layers across eight subjects and five folds for each model.

| Model | Median | Range |
| --- | --- | --- |
| Qwen2.5-VL-7B-Instruct | 89.73 | 87.90–93.25 |
| DeepSeek-VL-7B-Chat | 87.49 | 79.74–92.42 |
| LLaVA-v1.6-Mistral-7B | 93.84 | 88.53–96.13 |
| LLaVA-v1.6-Vicuna-7B | 92.42 | 88.89–97.12 |

**Figure A.1:**
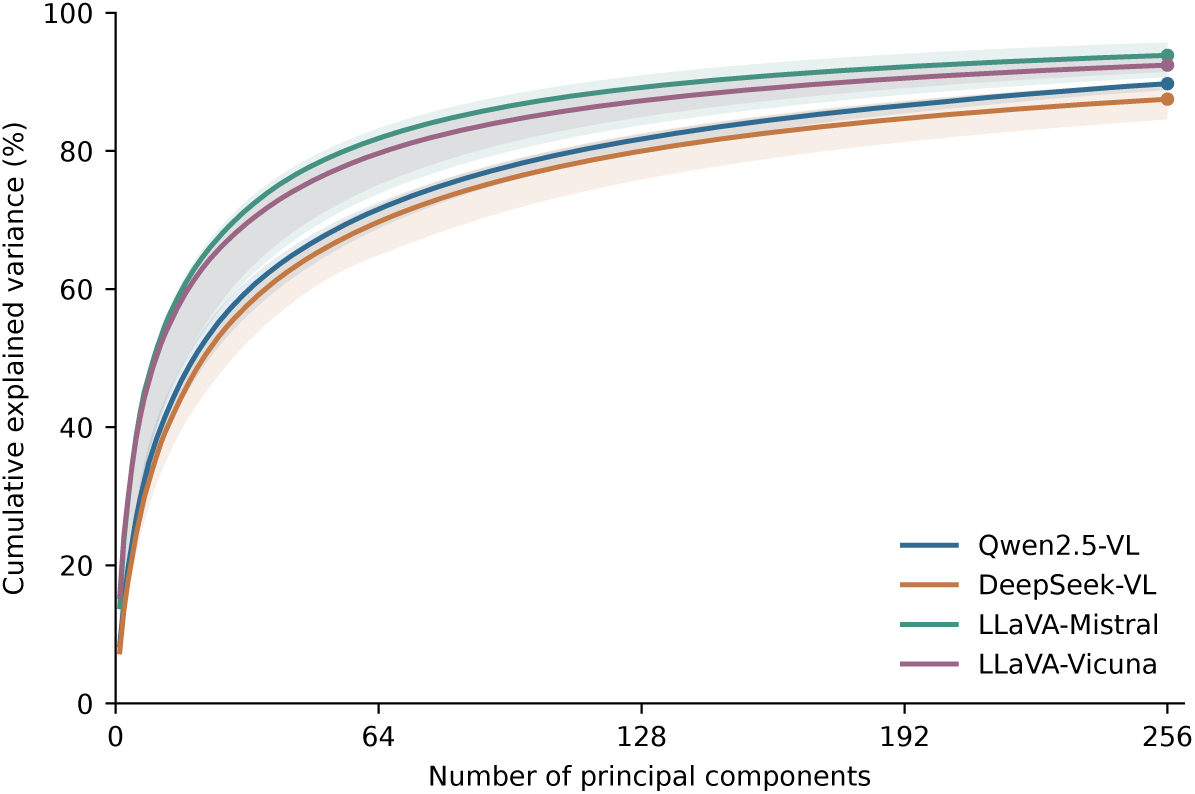
Cumulative explained variance over the first 256 principal components. Curves show the median training feature variance explained at each component count; shading denotes the interquartile range. Each model summarizes PCA fits for all selected language layers across eight subjects and five folds.

## B SEED Implementation

We compute pairwise image similarity using the released SEED implementation (Park et al., 2026), which combines Object F1, caption similarity, and EfficientNet feature similarity. All evaluation models remain frozen.

### Object agreement

We detect object categories with MM-Grounding-DINO (Zhao et al., 2024) using the category vocabulary provided by SEED. Let Prec(*p_a_, q_b_*) and Rec(*p_a_, q_b_*) denote the Object Precision and Object Recall computed following the released SEED implementation. Object agreement is their harmonic mean:

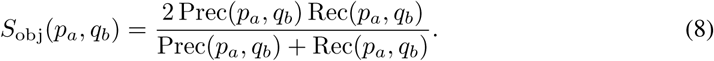

Object detection filtering and the aggregation of precision and recall follow the released SEED implementation.

### Caption similarity

We generate a caption for each image using GIT (Wang et al., 2022) and encode it with the Sentence Transformer (Reimers & Gurevych, 2019). Let **e**(*p*) denote the embedding of the caption generated for image *p*. Caption similarity is

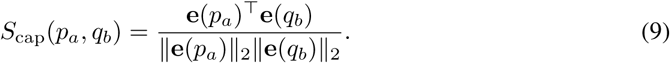

These captions are generated directly from the images; the human-written COCO captions used as MLLM inputs or for unit labeling are not supplied to the SEED evaluator.

### Visual similarity

We extract image representations using the pretrained EfficientNet model (Tan & Le, 2019) and preprocessing adopted by SEED. Let **f** (*p*) denote the resulting feature representation of image *p*. Following SEED, visual similarity is computed as the Pearson correlation between the two image representations:

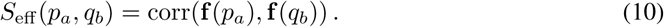

The final SEED score is the arithmetic mean of the three components, as defined in the main text. The same evaluation models and scoring procedure are used for all compared methods.

## C Preferred Image Collections and Optimal Transport

We construct both preferred image collections from the same held-out fold, with each collection containing *K* entries, following the procedure described in the main text. We treat each collection as an empirical distribution, assigning mass 1*/K* to each entry.

The feasible set in the main text is

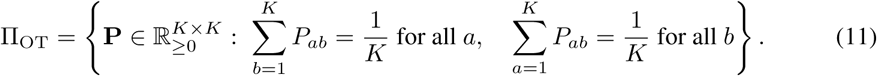

Here, *P_ab_* denotes the mass transported from the *a*-th entry in the voxel’s preferred image collection to the *b*-th entry in the SAE collection. The row and column constraints fix each entry’s outgoing and incoming mass, respectively, and ensure a total transported mass of 1.

## D Redundancy in SAE Unit Response Profiles Across Models

We extend the response-profile redundancy analysis to LLaVA-v1.6-Mistral-7B, LLaVA-v1.6-Vicuna-7B, and DeepSeek-VL-7B-Chat across eight subjects. The corresponding Qwen2.5-VL-7B-Instruct results are reported in the main text. Across the three additional models, SAIL exhibits lower redundancy than SAEs-BrainMap, with positive SAEs-BrainMap-minus-SAIL mean differences across the 200 resamples for every subject.

### D.1 Response-Profile Redundancy In LLAVA-V1.6-Mistral-7B

**Figure D.1:**
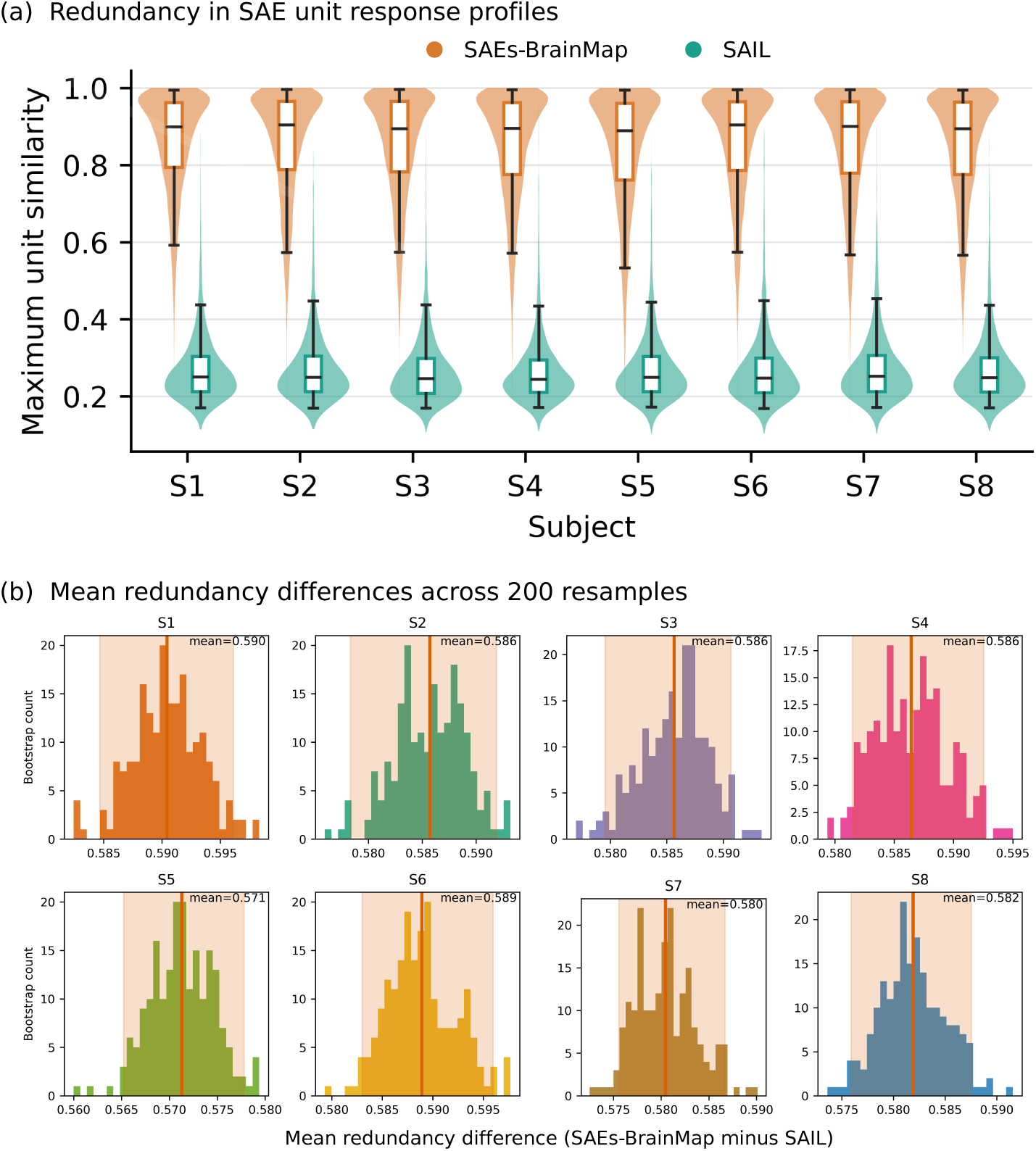
Unified cross-layer SAE training reduces unit response-profile redundancy in LLaVA-v1.6-Mistral-7B across subjects S1–S8. **(a)** Distributions of maximum cosine similarity between SAE unit response profiles for SAEs-BrainMap and SAIL across eight subjects. Both methods use matched numbers of active units and are evaluated on the same held-out images. **(b)** Distributions of the SAEs-BrainMap-minus-SAIL mean maximum unit similarity from 200 matched-size resamples. Lines denote means and shading denotes 95% intervals.

### D.2 Response-Profile Redundancy In Llava-V1.6-VICUNA-7B

**Figure D.2:**
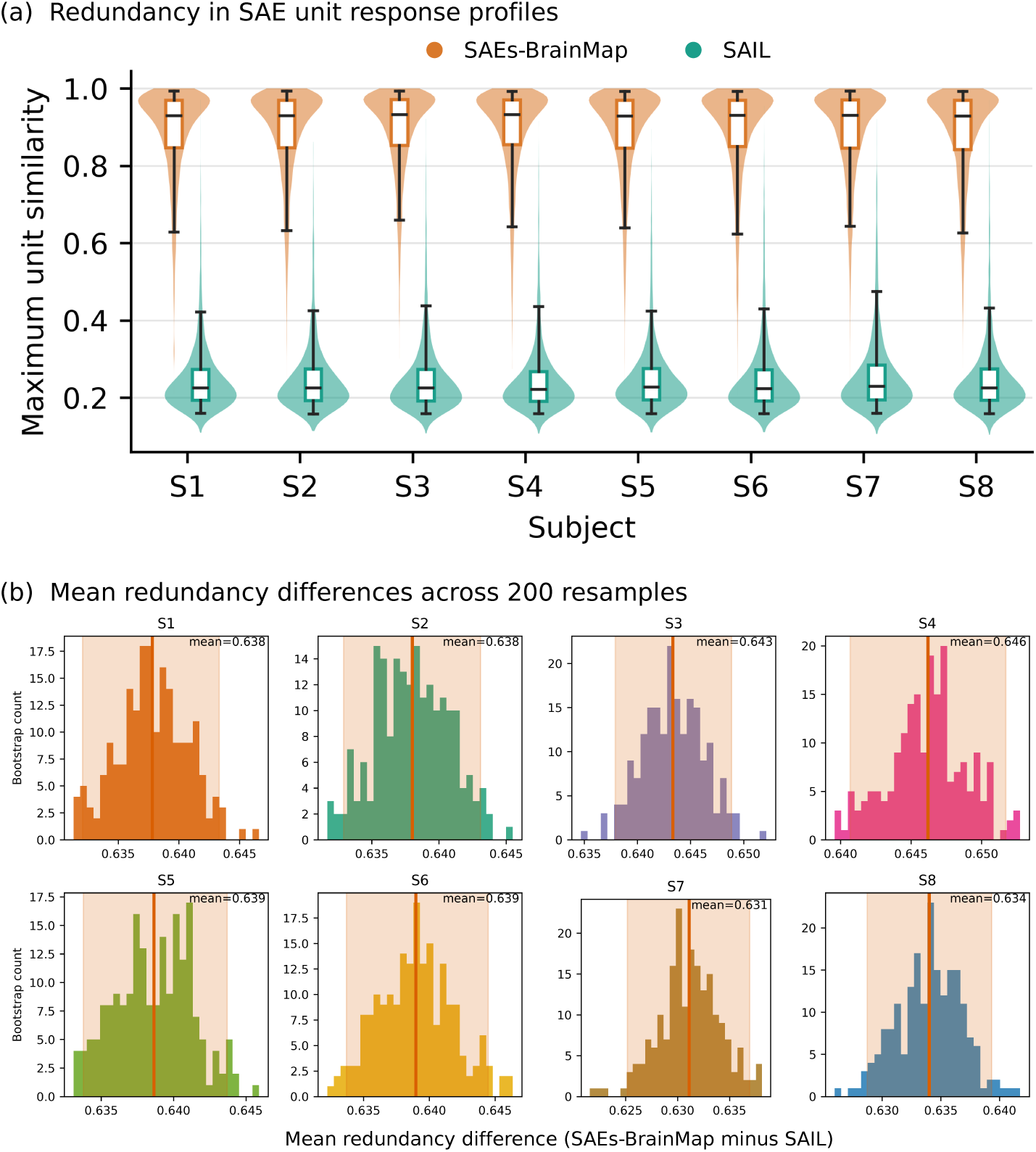
Unified cross-layer SAE training reduces unit response-profile redundancy in LLaVA-v1.6-Vicuna-7B across subjects S1–S8. **(a)** Distributions of maximum cosine similarity between SAE unit response profiles for SAEs-BrainMap and SAIL across eight subjects. Both methods use matched numbers of active units and are evaluated on the same held-out images. **(b)** Distributions of the SAEs-BrainMap-minus-SAIL mean maximum unit similarity from 200 matched-size resamples. Lines denote means and shading denotes 95% intervals.

### D.3 Response-Profile Redundancy in Deepseek-VL-7B-CHAT

**Figure D.3:**
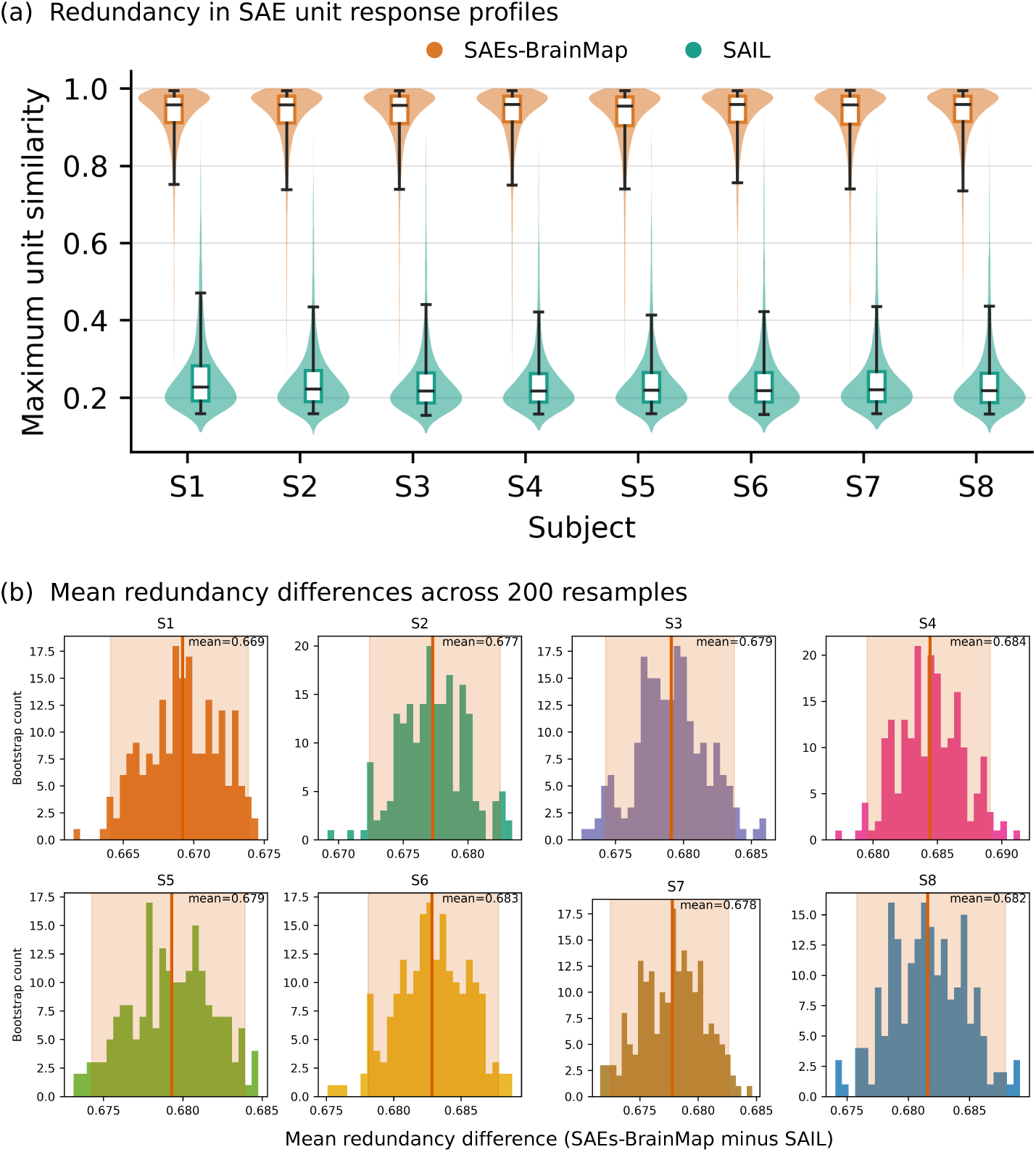
Unified cross-layer SAE training reduces unit response-profile redundancy in DeepSeek-VL-7B-Chat across subjects S1–S8. **(a)** Distributions of maximum cosine similarity between SAE unit response profiles for SAEs-BrainMap and SAIL across eight subjects. Both methods use matched numbers of active units and are evaluated on the same held-out images. **(b)** Distributions of the SAEs-BrainMap-minus-SAIL mean maximum unit similarity from 200 matched-size resamples. Lines denote means and shading denotes 95% intervals.

## E Semantic Alignment Comparisons Across Models

The following plots summarize semantic alignment for the two LLaVA models and DeepSeek; the Qwen comparison appears in the main text. Upper and lower panels show Selective ROI and Whole cortex, respectively. Each point represents one subject, and each short horizontal line marks the arithmetic mean across the eight subjects.

**Figure E.1:**
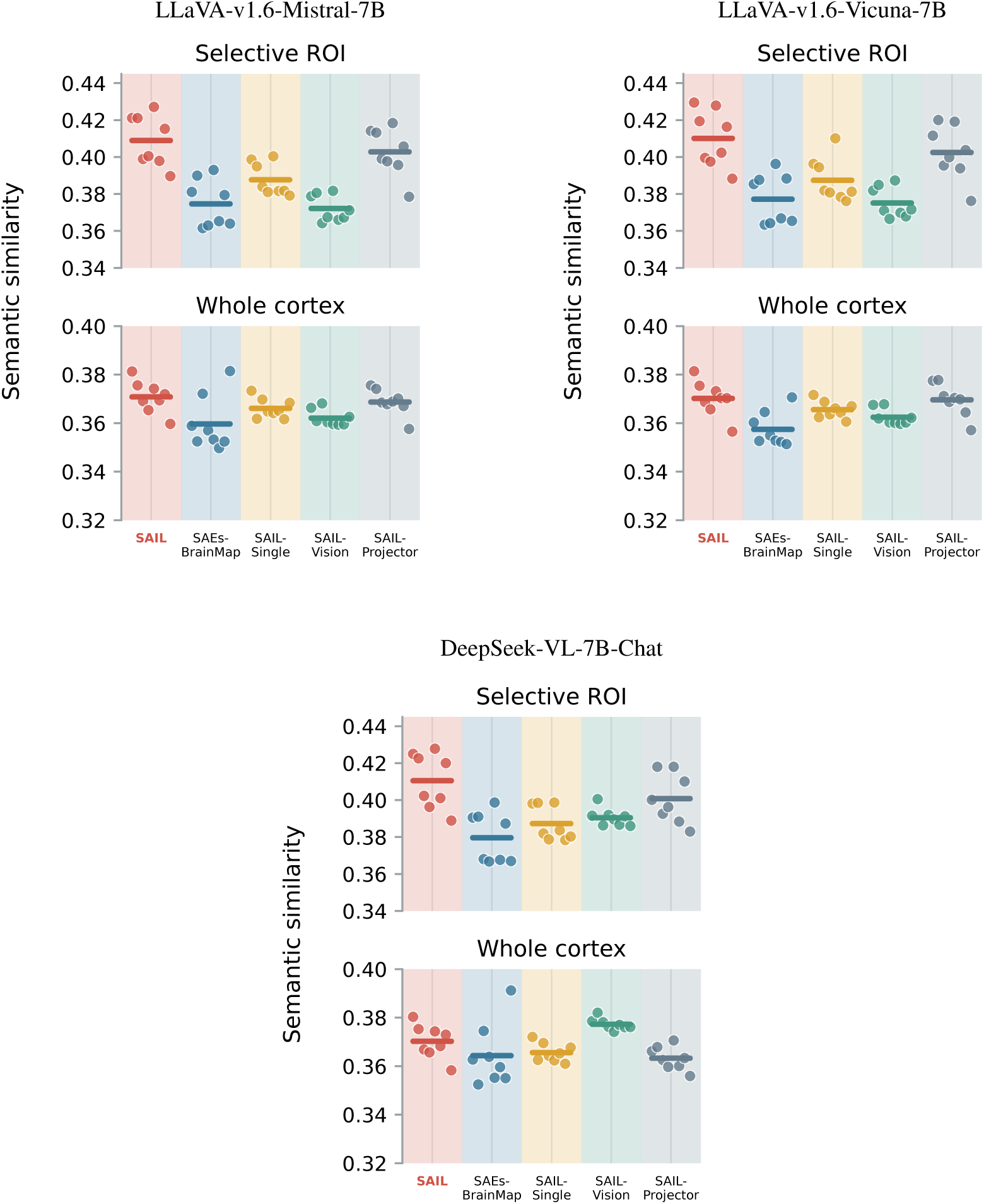
Semantic alignment summaries across eight subjects for each model.

## F Searchlight RSA: Evaluation Settings and Full Results

Searchlight RSA evaluates whether model representations capture the pattern of brain response differences across images, complementing the semantic comparison of preferred image collections. We report results for all four models and eight subjects. The primary comparison is between SAIL and SAEs-BrainMap; RAW, PCA, and ICA provide additional reference representations.

Each searchlight comprises at least 20 cortical voxels within a radius of 3 in voxel-grid coordinates. We construct the brain RDM from correlation distances between local response vectors.

For RAW, PCA, and ICA, we compute comparison RDMs directly from image features. For SAIL and SAEs-BrainMap, we use the previously obtained voxel-unit correspondences to predict held-out voxel responses and construct predicted-response RDMs within the same cortical neighborhoods.

For each searchlight, we correlate the upper triangles of the brain and comparison RDMs using Pearson correlation. Tables summarize valid searchlights within the Whole cortex and Selective ROI using Mean, Median, and Top 5% mean. The following tables and maps report these comparisons separately for each model and subject, with bold entries indicating column maxima (including ties at the reported precision).

### F.1 Searchlight RSA of Qwen2.5-VL-7B-Instruct Representations

**Table F.1:**
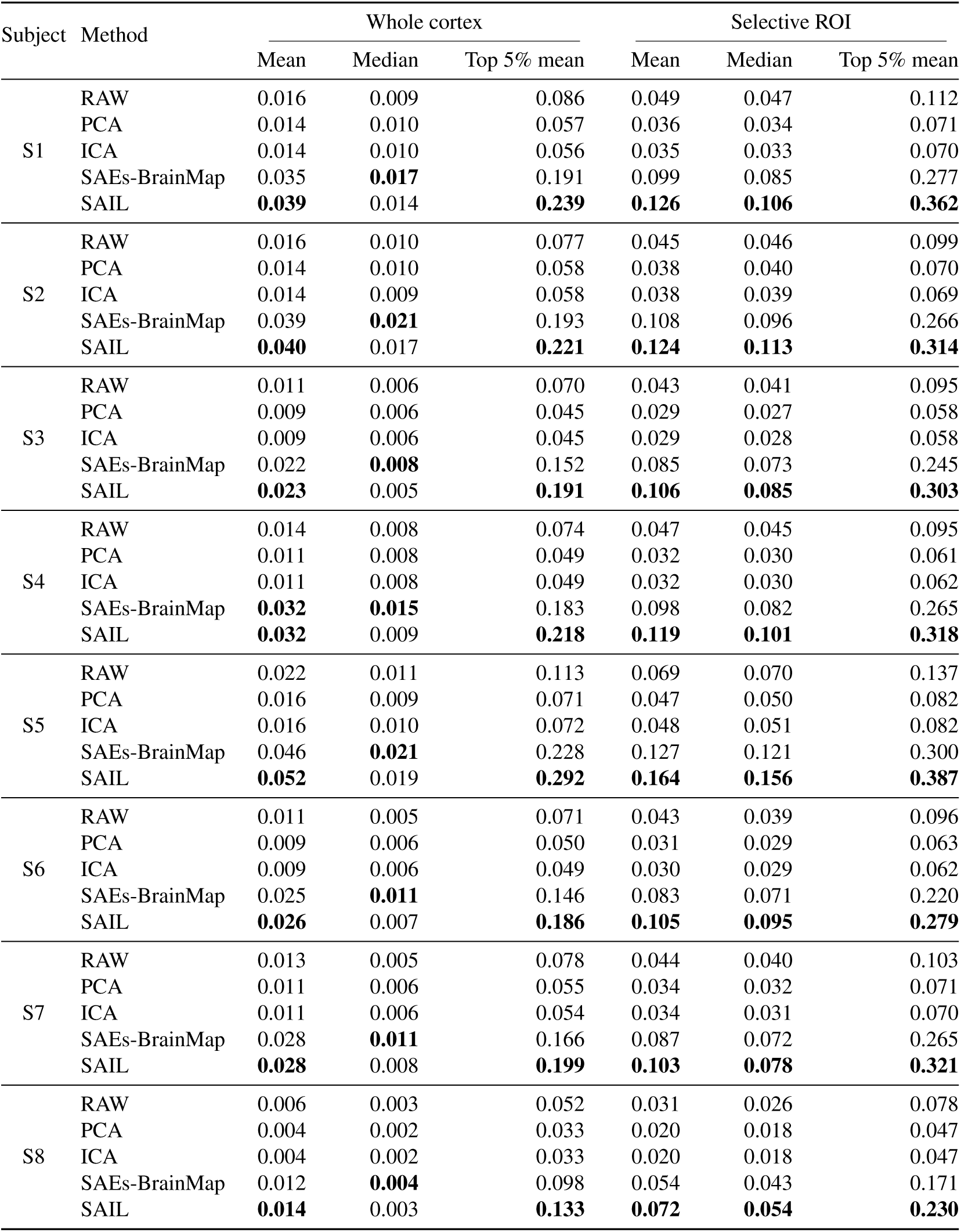
Searchlight RSA across five representation methods for Qwen2.5-VL-7B-Instruct in subjects S1–S8. Bold entries indicate the highest value within each subject and metric, including ties at the reported precision.

**Figure F.1:**
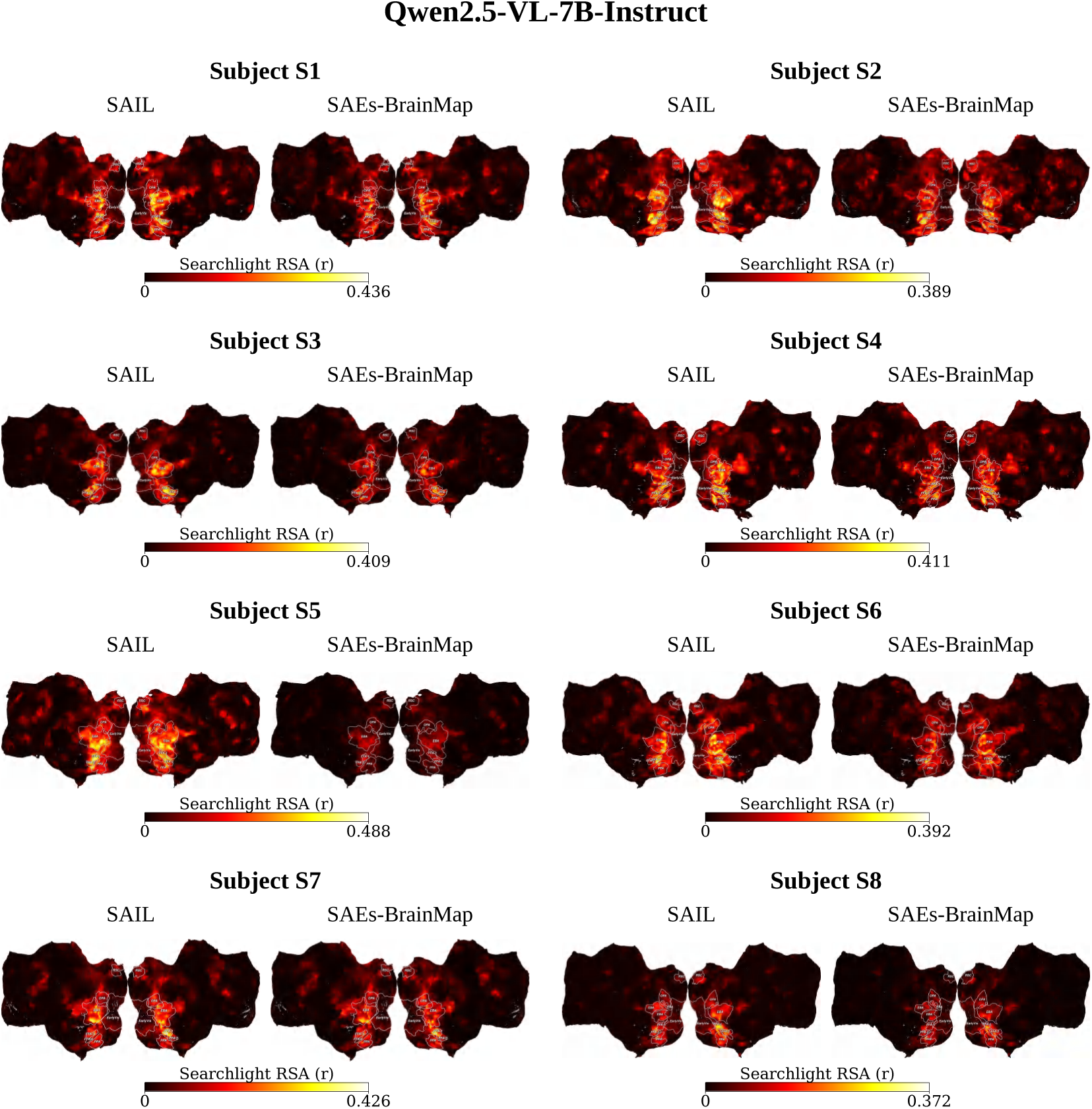
Cortical distribution of searchlight RSA scores for Qwen2.5-VL-7B-Instruct in subjects S1–S8, ordered from left to right and top to bottom. Each subject panel compares SAIL (left) with SAEs-BrainMap (right) using the same subject-specific color scale.

### F.2 Searchlight RSA of LLaVA-v1.6-Mistral-7B Representations

**Table F.2:**
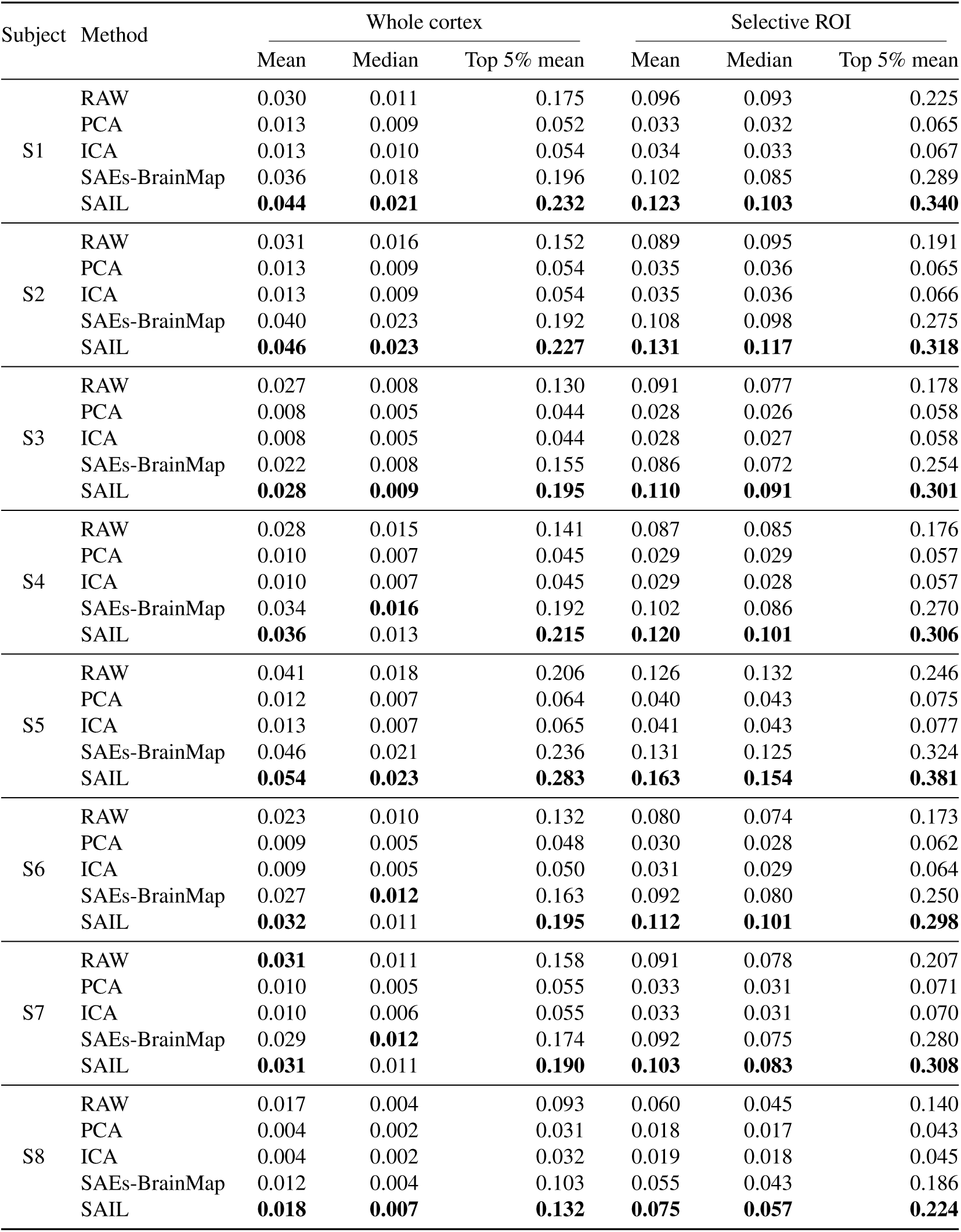
Searchlight RSA across five representation methods for LLaVA-v1.6-Mistral-7B in subjects S1–S8. Bold entries indicate the highest value within each subject and metric, including ties at the reported precision.

**Figure F.2:**
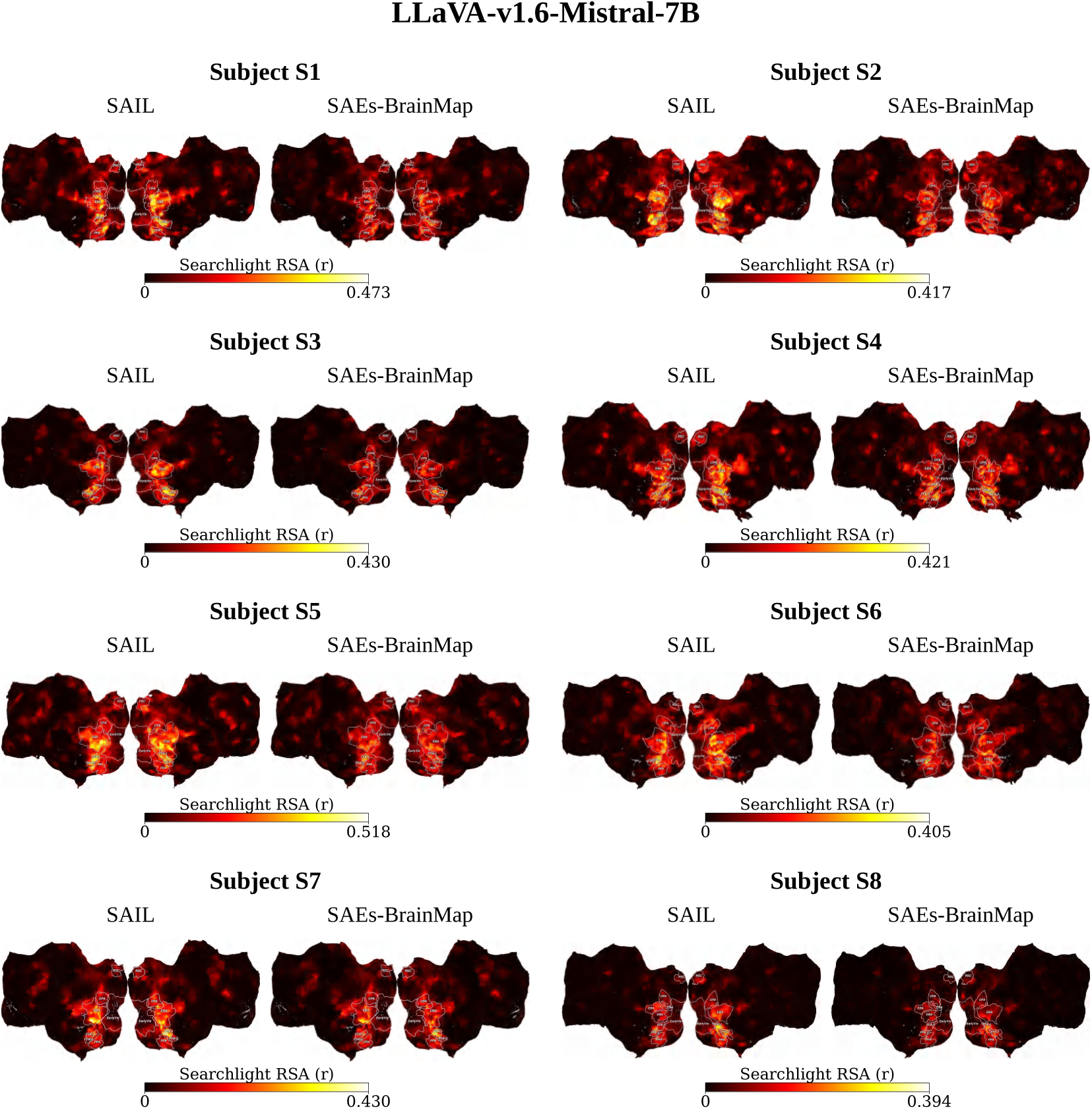
Cortical distribution of searchlight RSA scores for LLaVA-v1.6-Mistral-7B in subjects S1–S8, ordered from left to right and top to bottom. Each subject panel compares SAIL (left) with SAEs-BrainMap (right) using the same subject-specific color scale.

### F.3 Searchlight RSA of LLaVA-v1.6-Vicuna-7B Representations

**Table F.3:**
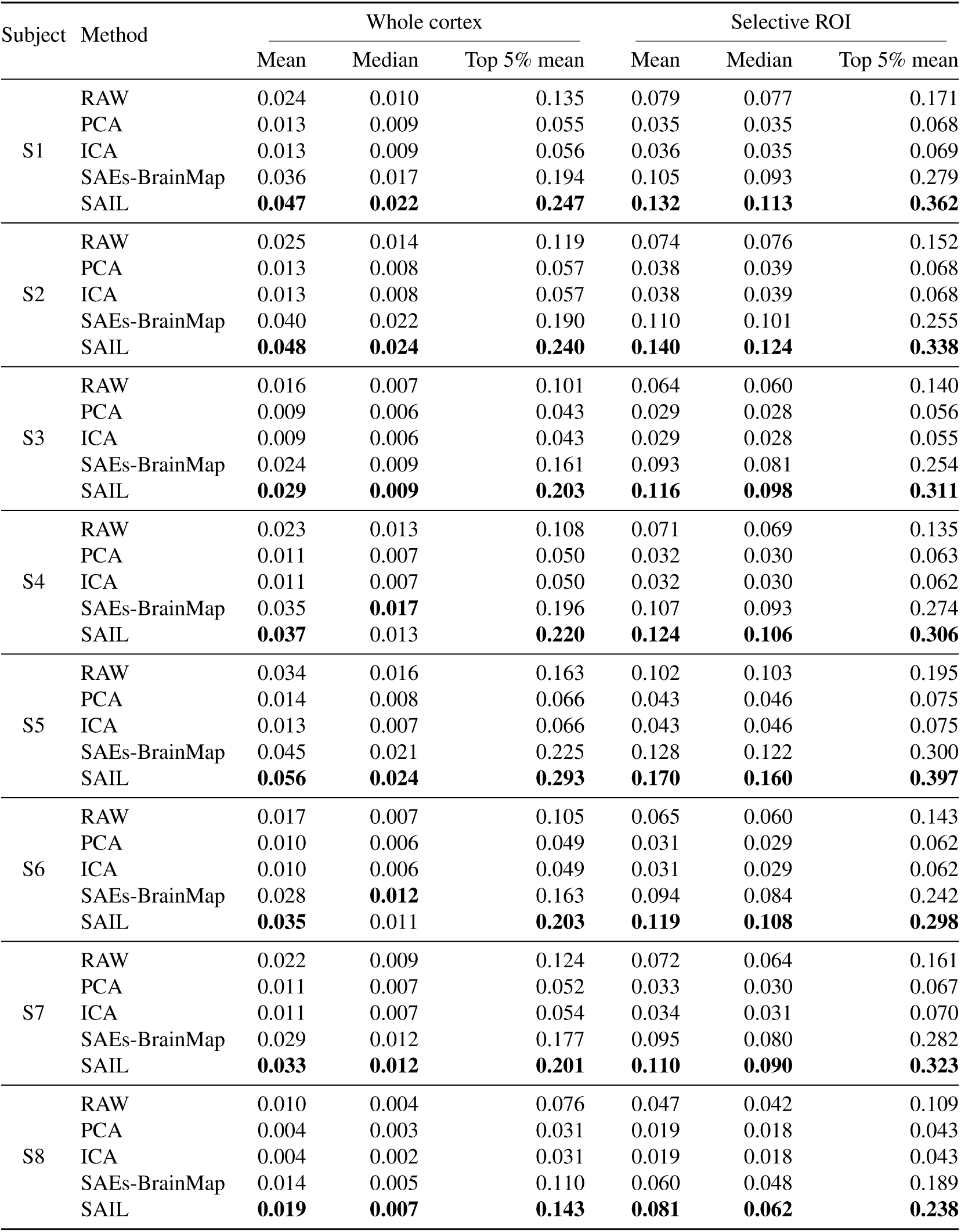
Searchlight RSA across five representation methods for LLaVA-v1.6-Vicuna-7B in subjects S1–S8. Bold entries indicate the highest value within each subject and metric, including ties at the reported precision.

**Figure F.3:**
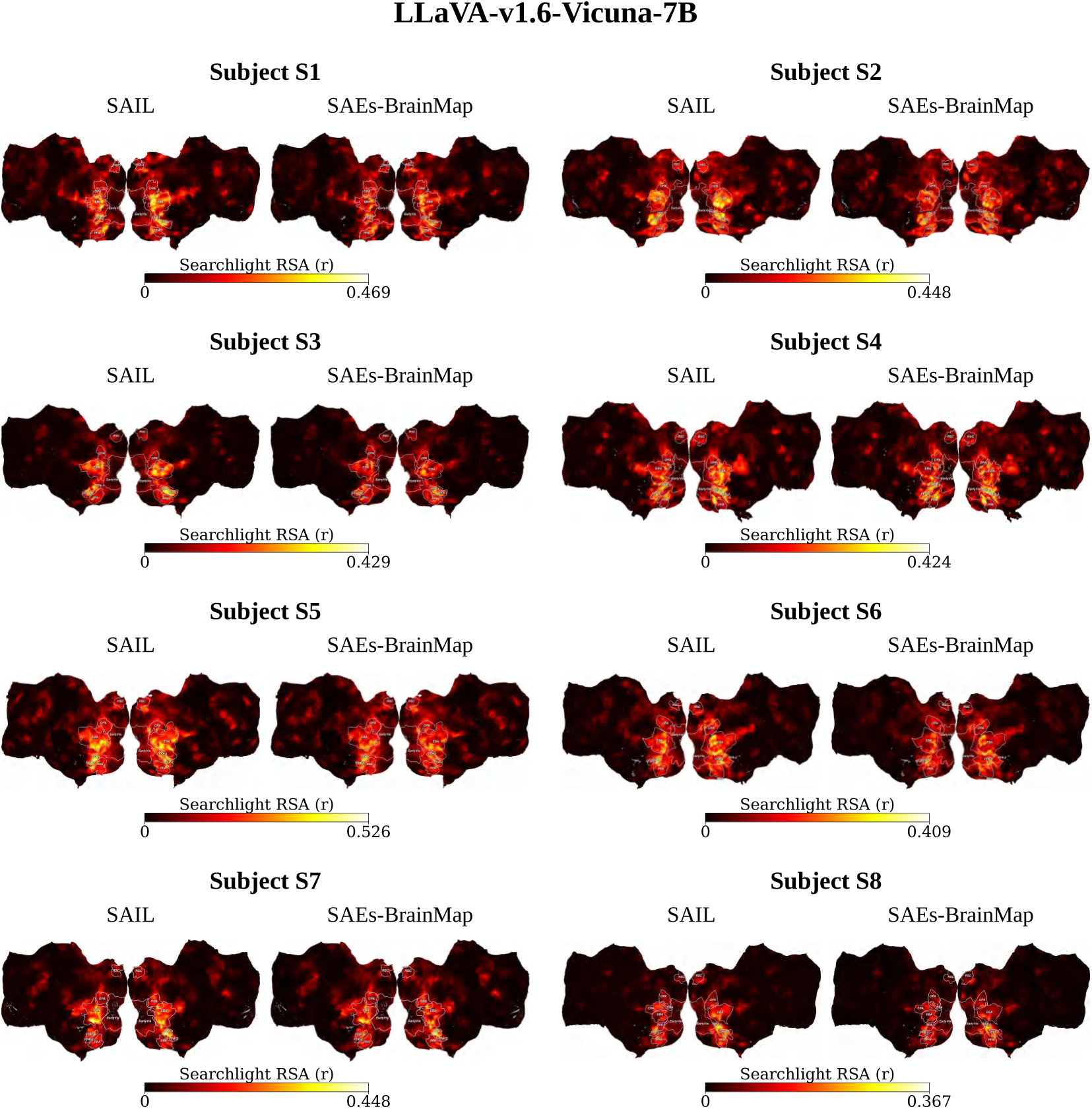
Cortical distribution of searchlight RSA scores for LLaVA-v1.6-Vicuna-7B in subjects S1–S8, ordered from left to right and top to bottom. Each subject panel compares SAIL (left) with SAEs-BrainMap (right) using the same subject-specific color scale.

### F.4 Searchlight RSA of DeepSeek-VL-7B-Chat Representations

**Table F.4:**
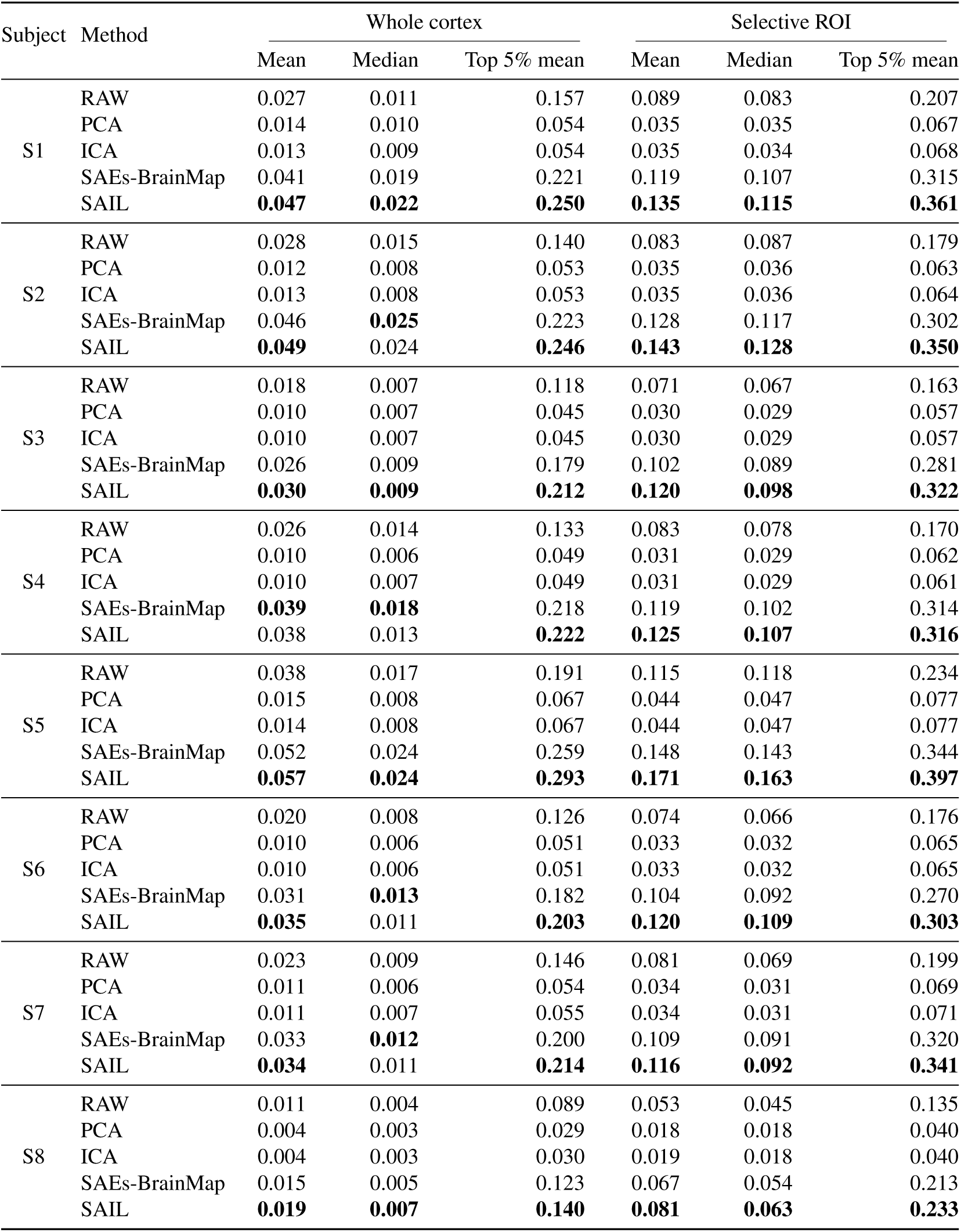
Searchlight RSA across five representation methods for DeepSeek-VL-7B-Chat in subjects S1–S8. Bold entries indicate the highest value within each subject and metric, including ties at the reported precision.

**Figure F.4:**
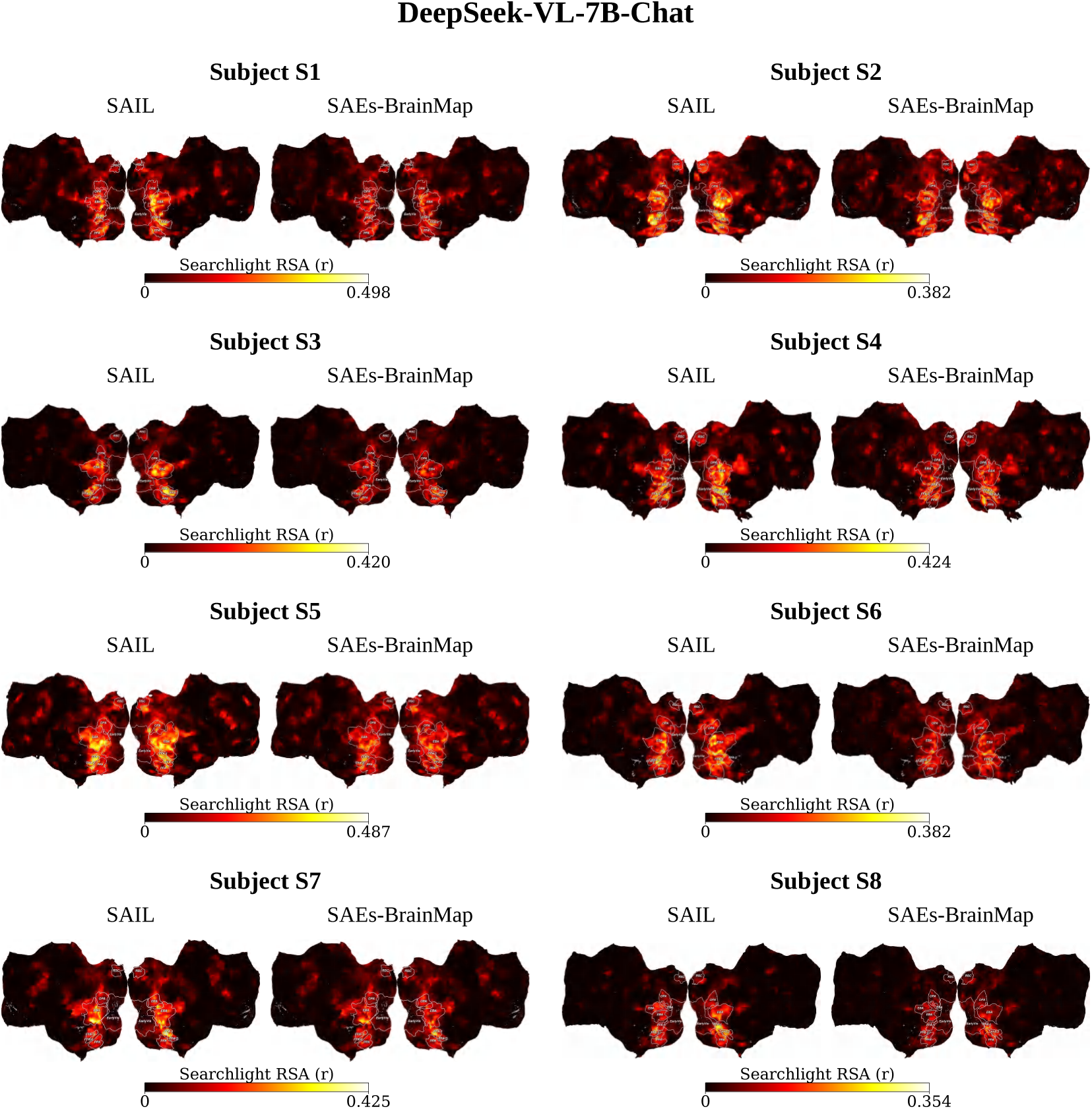
Cortical distribution of searchlight RSA scores for DeepSeek-VL-7B-Chat in subjects S1–S8, ordered from left to right and top to bottom. Each subject panel compares SAIL (left) with SAEs-BrainMap (right) using the same subject-specific color scale.

### F.5 RSA with Matched Candidate Dimensionality

To assess whether our results are sensitive to differences in candidate dimensionality, we subsample active SAE units without replacement so that the number of SAE features matches the number of features in the corresponding raw model representations (RAW). We repeat this resampling procedure 100 times for each of four models and eight subjects, and summarize the resulting distributions in the figures.

F.5.1 RSA with Matched Dimensionality for Qwen2.5-VL-7B-Instruct

**Figure F.5:**
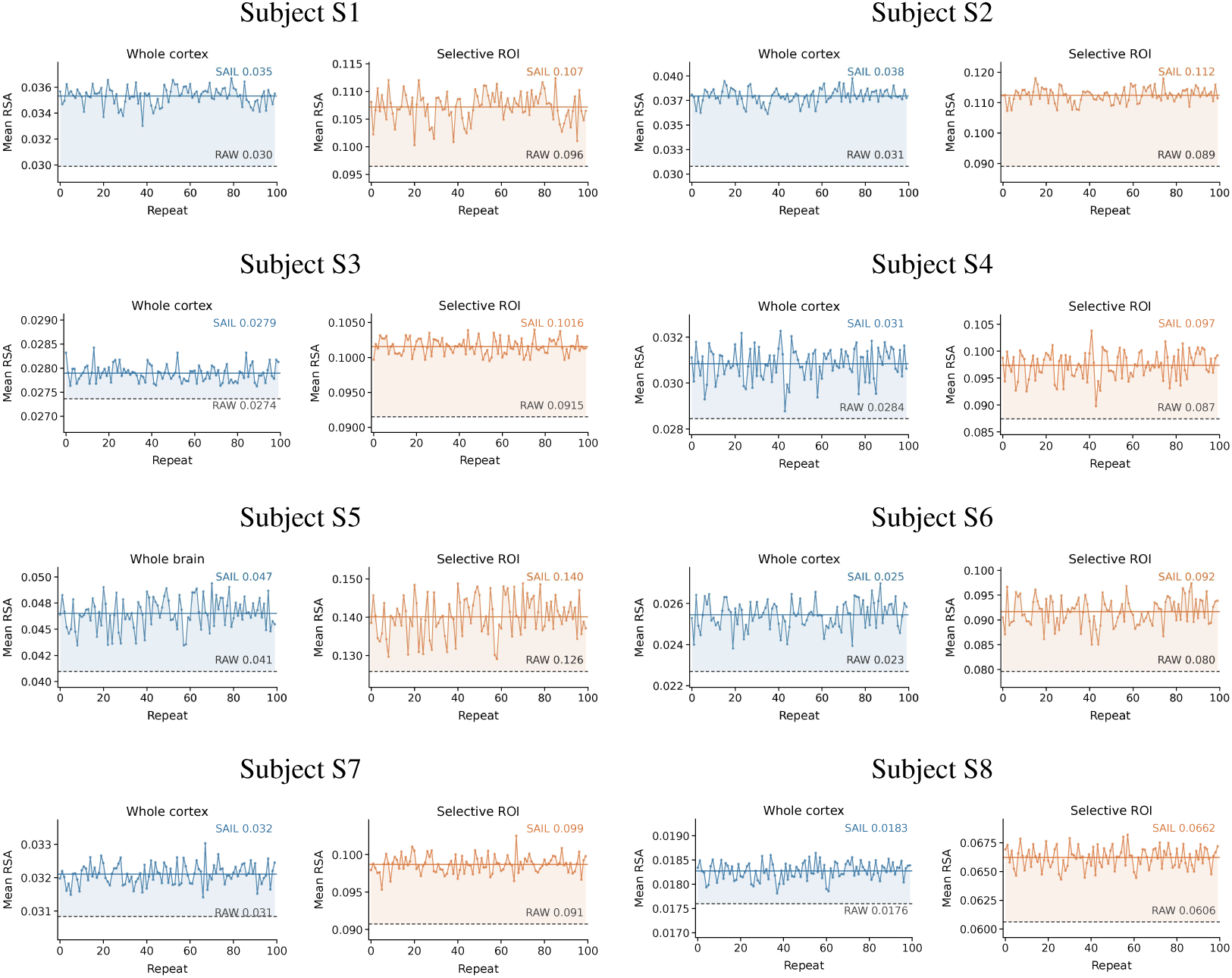
RSA with matched candidate dimensionality for Qwen2.5-VL-7B-Instruct across subjects S1–S8. Each row contains two subjects. Within each subject, the left and right plots show mean searchlight RSA over the Whole cortex and within the Selective ROI, respectively. Colored curves show SAIL scores across 100 unit-resampling repeats; solid horizontal lines indicate their means. RAW denotes the original MLLM representations from the final language layer, without PCA, ICA, or SAE decomposition.

F.5.2 RSA with Matched Dimensionality for LLaVA-v1.6-Mistral-7B Representations

**Figure F.6:**
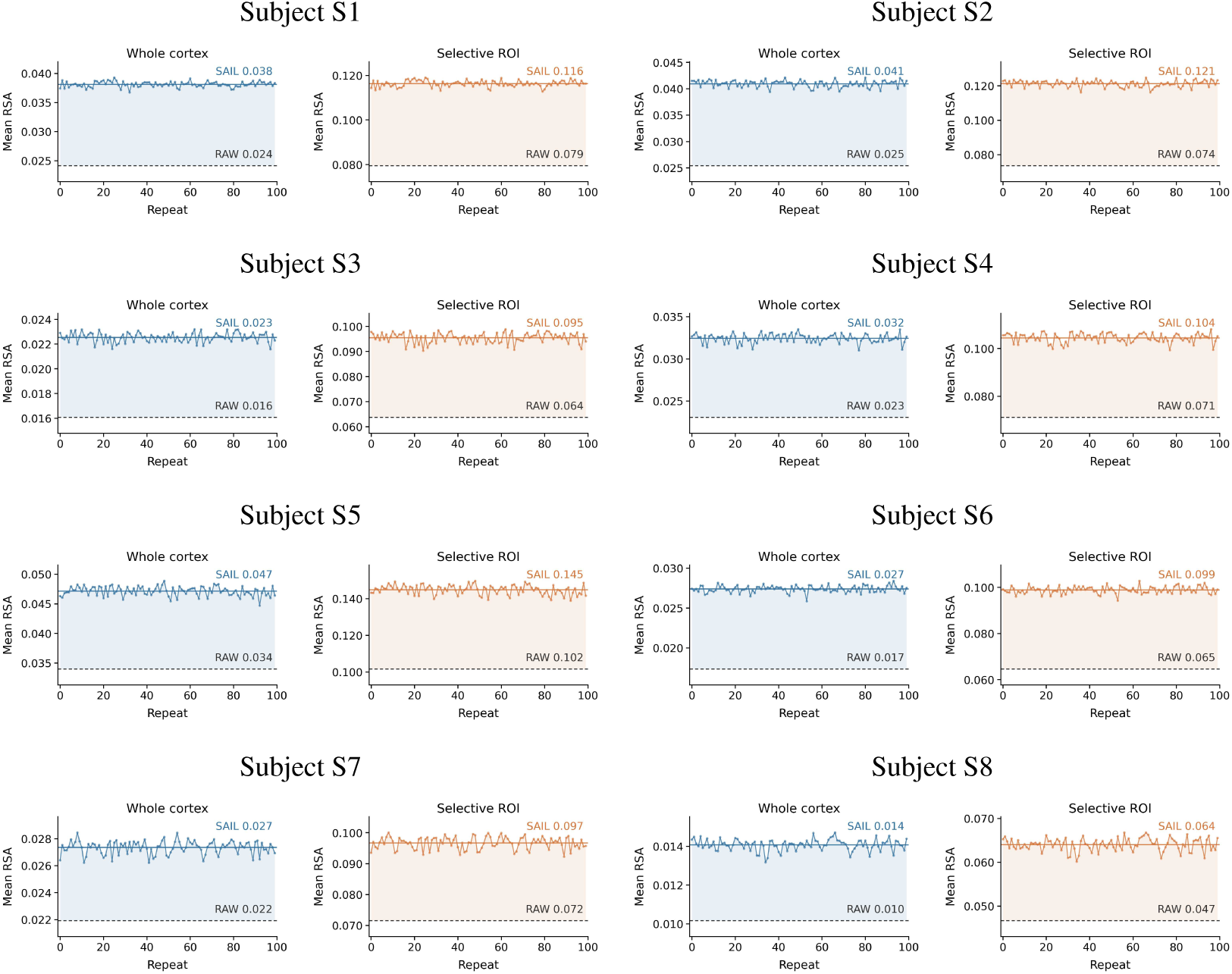
RSA with matched candidate dimensionality for LLaVA-v1.6-Mistral-7B across subjects S1–S8. Each row contains two subjects. Within each subject, the left and right plots show mean searchlight RSA over the Whole cortex and within the Selective ROI, respectively. Colored curves show SAIL scores across 100 unit-resampling repeats; solid horizontal lines indicate their means. RAW denotes the original MLLM representations from the final language layer, without PCA, ICA, or SAE decomposition.

F.5.3 RSA with Matched Dimensionality for LLaVA-v1.6-Vicuna-7B Representations

**Figure F.7:**
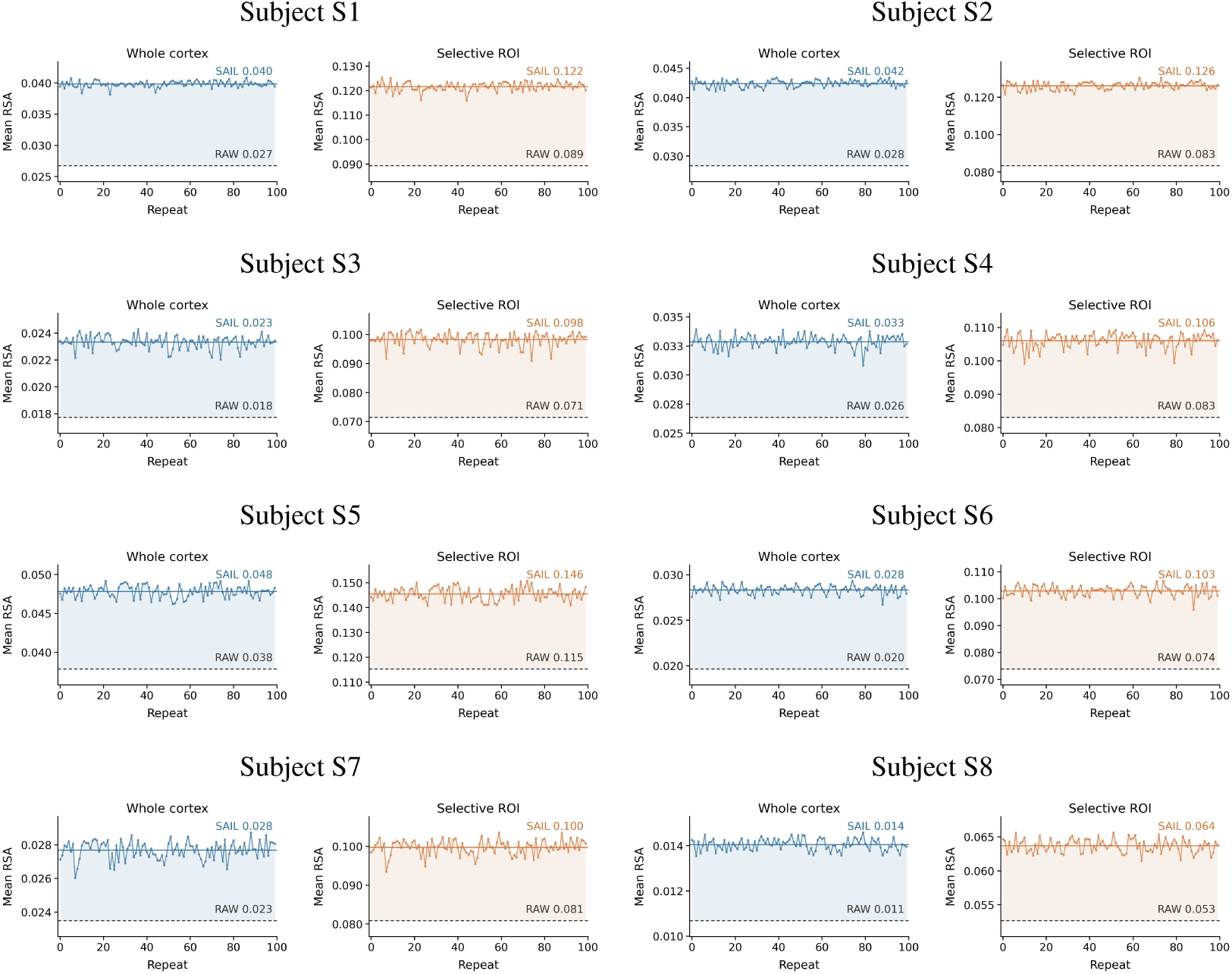
RSA with matched candidate dimensionality for LLaVA-v1.6-Vicuna-7B across subjects S1–S8. Each row contains two subjects. Within each subject, the left and right plots show mean searchlight RSA over the Whole cortex and within the Selective ROI, respectively. Colored curves show SAIL scores across 100 unit-resampling repeats; solid horizontal lines indicate their means. RAW denotes the original MLLM representations from the final language layer, without PCA, ICA, or SAE decomposition.

F.5.4 RSA with Matched Dimensionality for DeepSeek-VL-7B-Chat

**Figure F.8:**
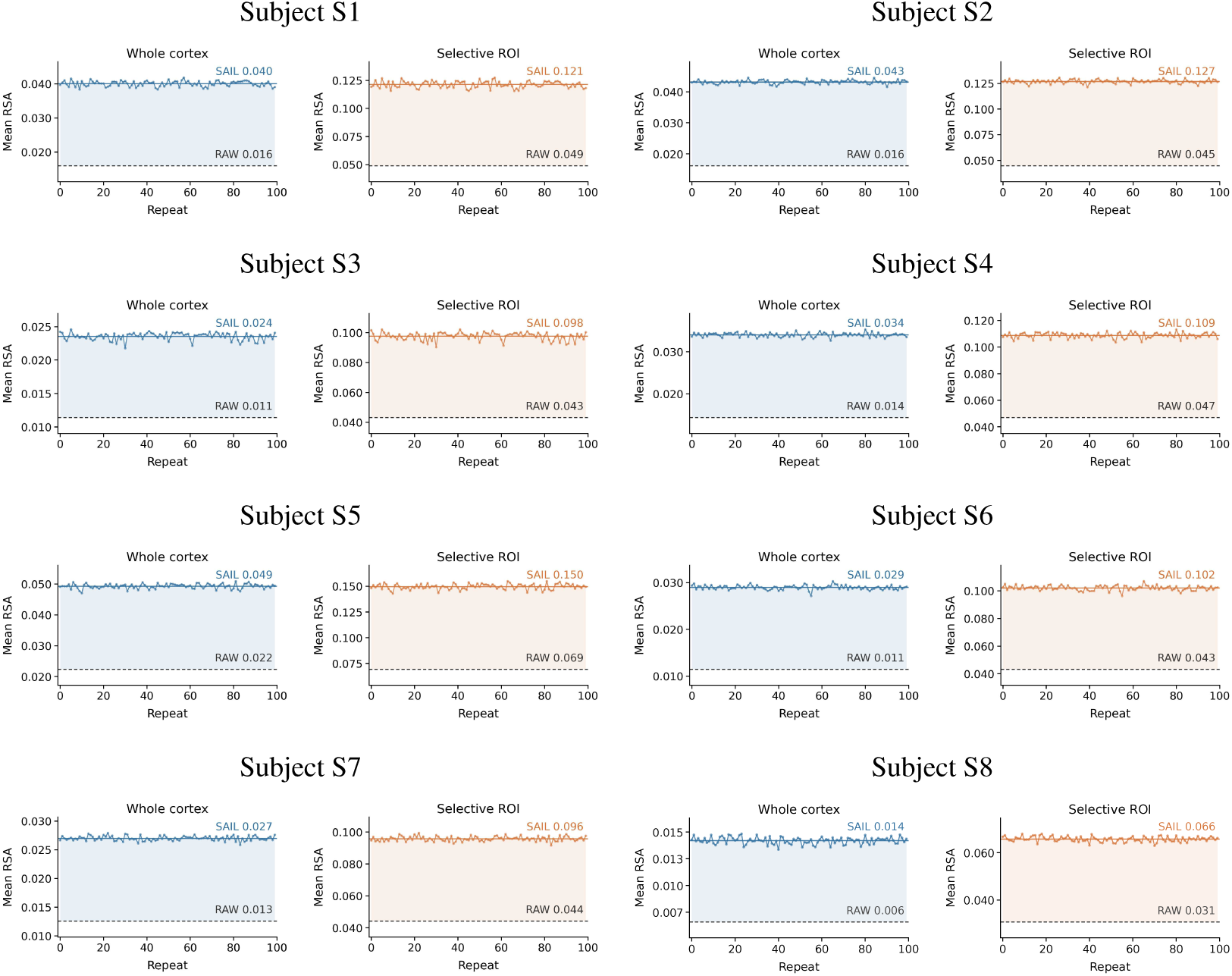
RSA with matched candidate dimensionality for DeepSeek-VL-7B-Chat across subjects S1–S8. Each row contains two subjects. Within each subject, the left and right plots show mean searchlight RSA over the Whole cortex and within the Selective ROI, respectively. Colored curves show SAIL scores across 100 unit-resampling repeats; solid horizontal lines indicate their means. RAW denotes the original MLLM representations from the final language layer, without PCA, ICA, or SAE decomposition.

## G Cortical Semantic Maps Across Models and Subjects

We extend the cortical semantic visualization to four models and eight subjects; the Qwen map for subject S5 appears in the main text and is omitted here.

### G.1 QWEN2.5-VL-7B-INSTRUCT

**Figure G.1:**
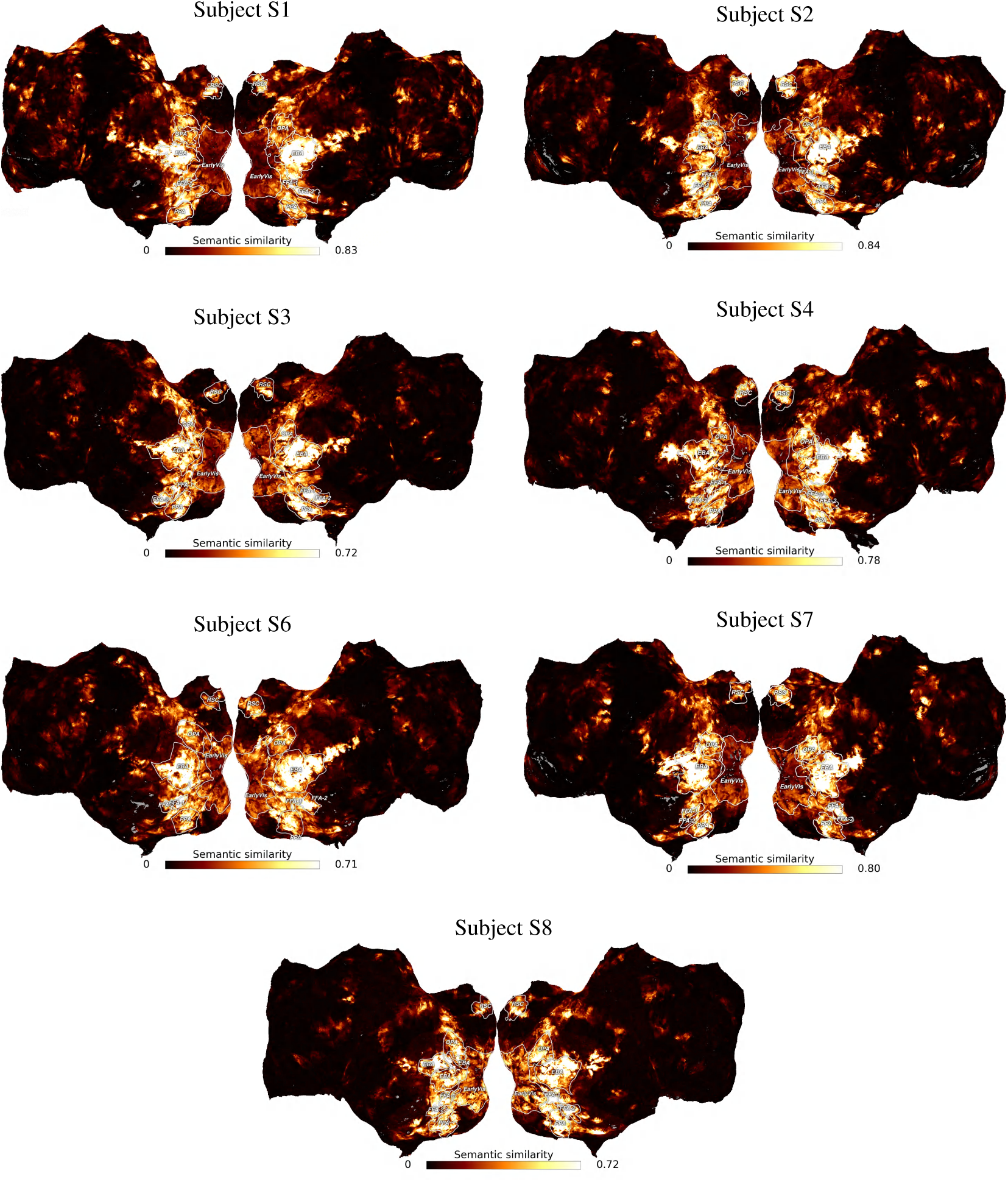
Cortical semantic maps for Qwen2.5-VL-7B-Instruct in subjects S1–S4 and S6–S8.

### G.2 LLAVA-V1.6-MISTRAL-7B

**Figure G.2:**
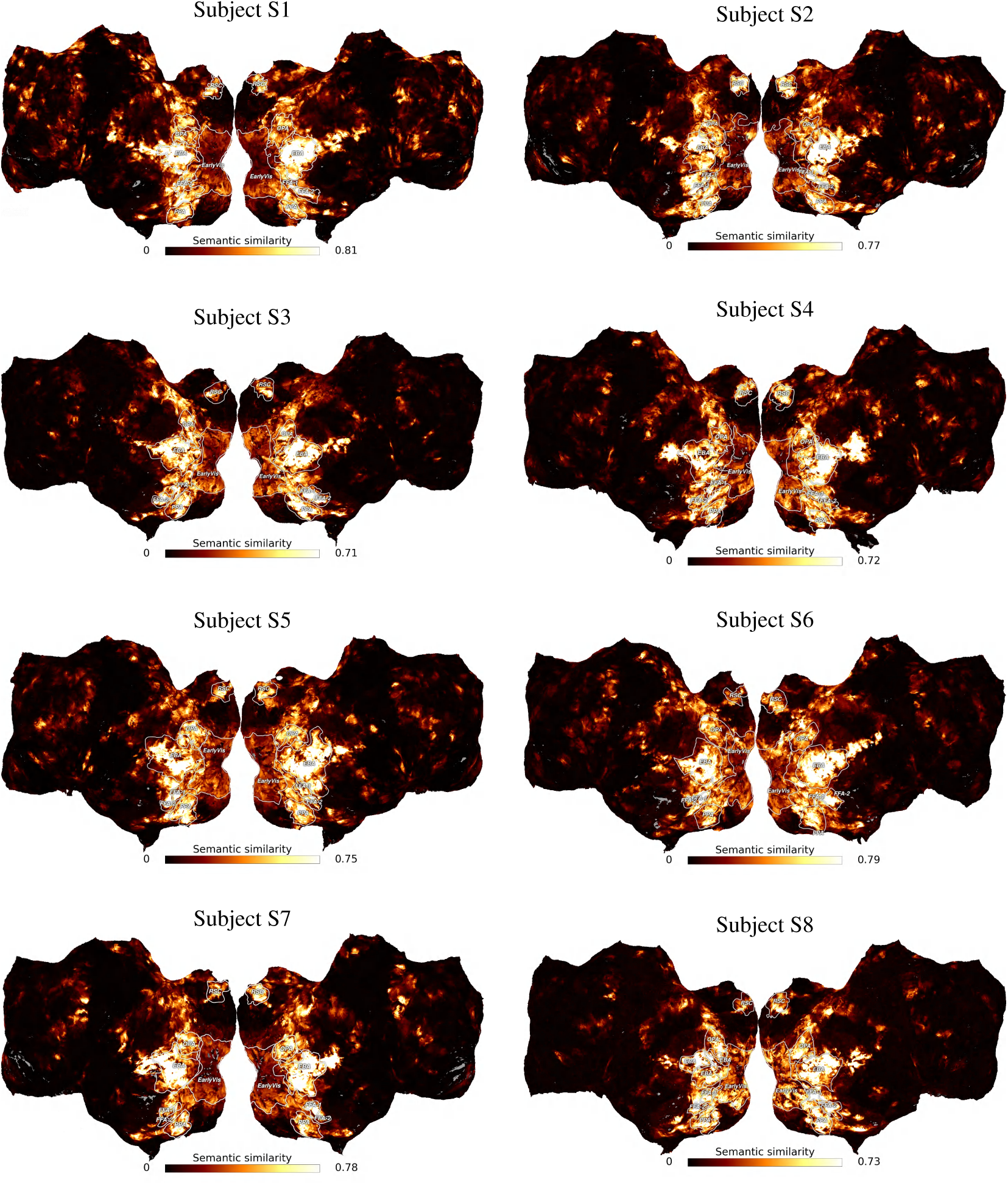
Cortical semantic maps for LLaVA-v1.6-Mistral-7B in subjects S1–S8.

### G.3 LLAVA-V1.6-VICUNA-7B

**Figure G.3:**
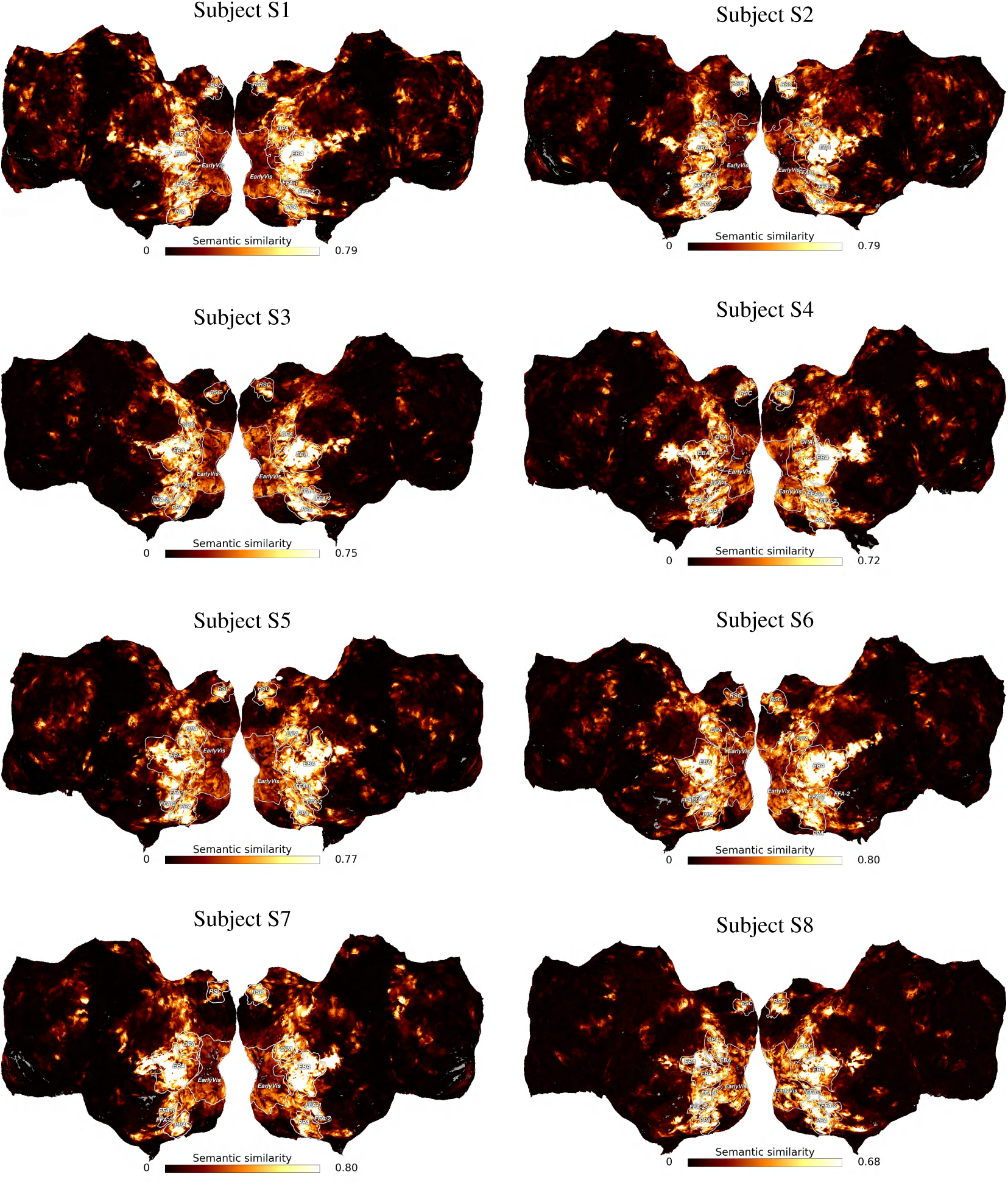
Cortical semantic maps for LLaVA-v1.6-Vicuna-7B in subjects S1–S8.

### G.4 DEEPSEEK-VL-7B-CHAT

**Figure G.4:**
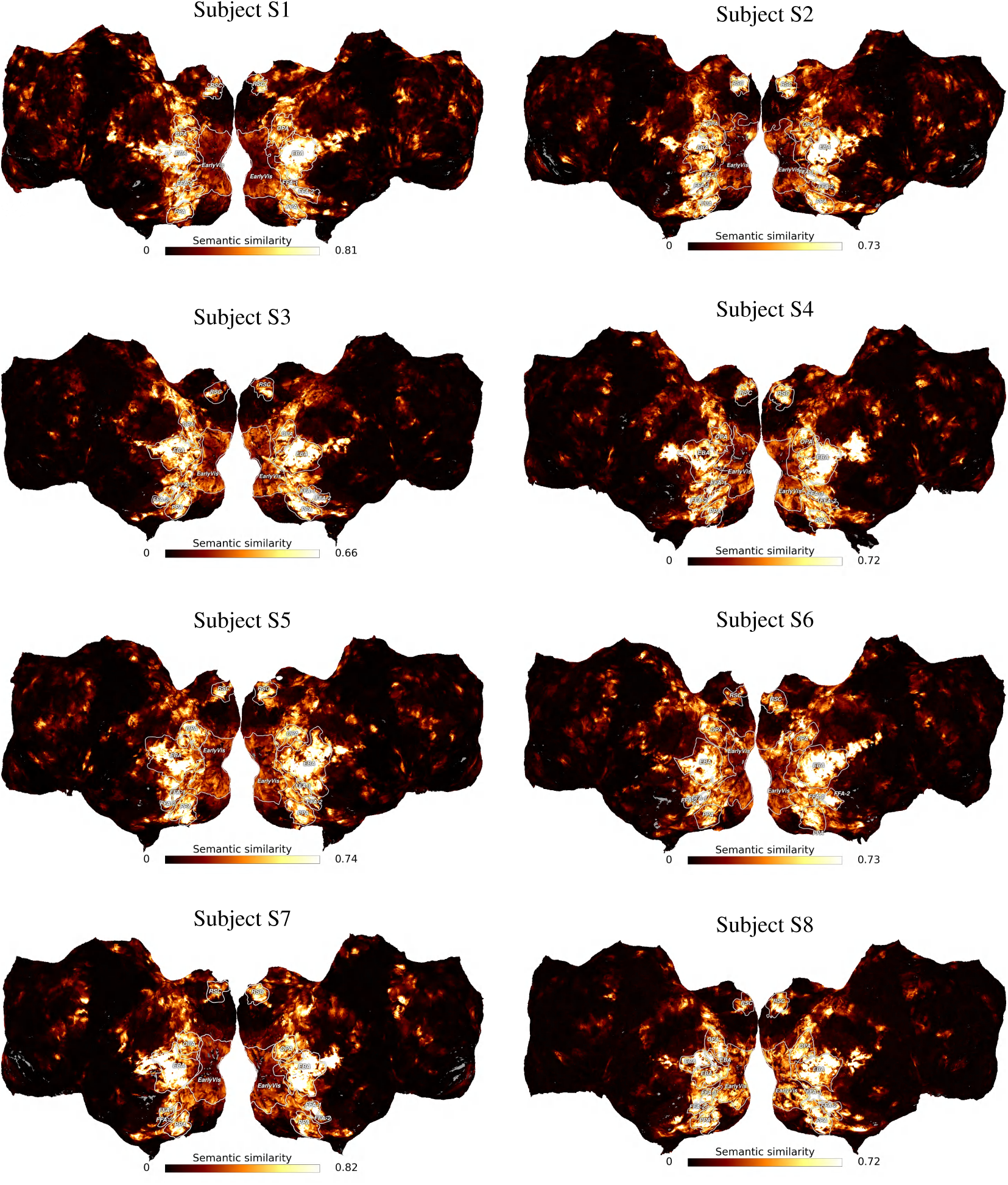
Cortical semantic maps for DeepSeek-VL-7B-Chat in subjects S1–S8.

## H Semantic Label Assignment

For each matched SAE unit, we use its maximally activating image from the corresponding held-out image pool and retrieve the five associated COCO captions. Candidate labels are extracted automatically by lowercasing, tokenizing, and lemmatizing the captions. We retain nouns, verbs, and adjectives, remove English stopwords and generic image-description terms, and deduplicate the resulting candidates. Multi-word noun phrases are retained when they have a noun entry in WordNet. To select the label, we keep the image fixed and insert each candidate into the same text template, “a photo of *{*candidate*}*”. Each image-text pair is processed by the frozen MLLM, followed by the same sequence pooling, layer-wise PCA, standardization, and layer concatenation used in the main analysis. The resulting representation is passed through the corresponding frozen SAE encoder, and we record the activation of the target unit. All fitted transformations and model parameters remain unchanged during this procedure. The candidate eliciting the highest target-unit activation is assigned as the semantic label. This activation is computed from the pooled representation of the complete image–text input. Candidate extraction and ranking do not use fMRI responses, ROI membership, or cortical location. Each voxel receives the labels assigned to its matched SAE units.

## I fLoc Category Response Reconstruction Across Models

We report fLoc category response reconstruction for the two LLaVA models and DeepSeek across eight subjects, complementing the Qwen results in the main text. The tables give cosine similarity between predicted and measured response vectors for each category and subject.

**Table I.1:**
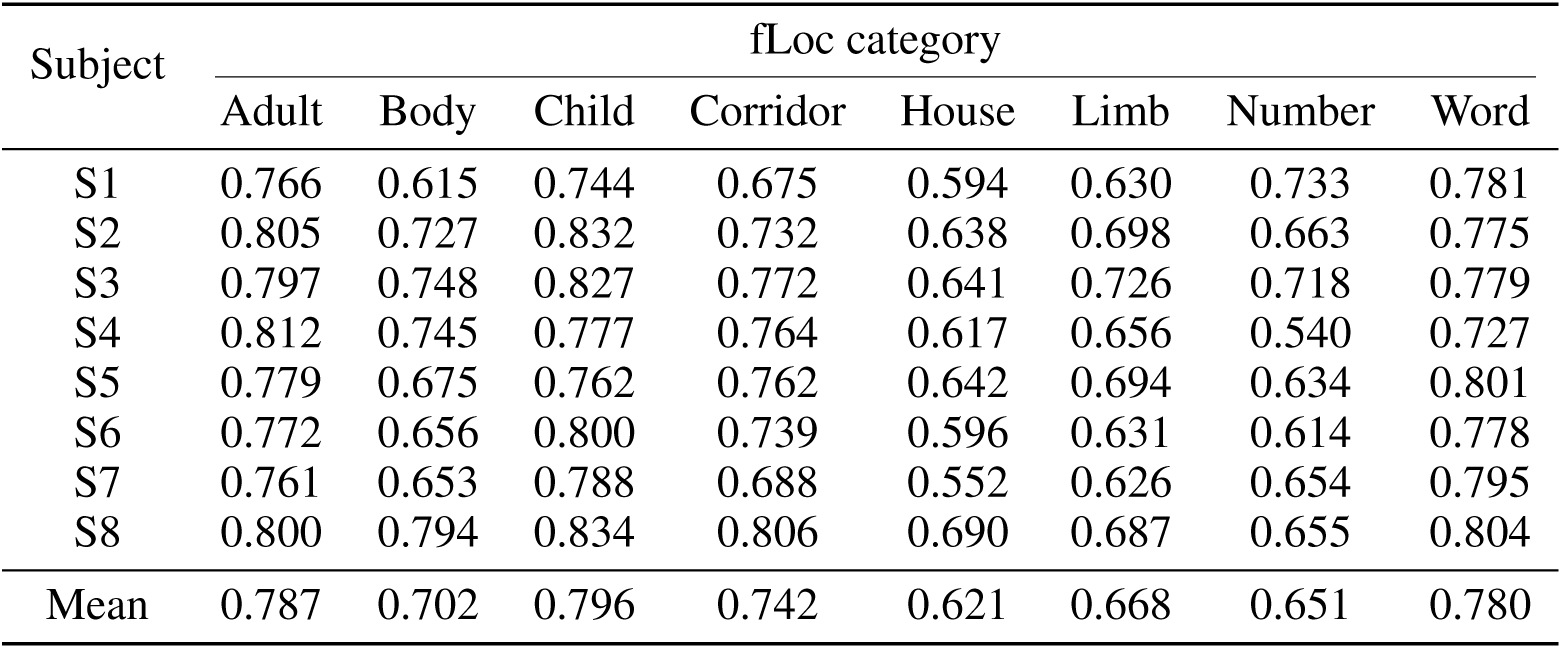
Cosine similarity between SAIL predicted and measured fLoc category responses for LLaVA-v1.6-Mistral-7B.

**Table I.2:** Cosine similarity between SAIL predicted and measured fLoc category responses for LLaVA-v1.6-Vicuna-7B.

| Subject | fLoc category |  |  |  |  |  |  |  |
| --- | --- | --- | --- | --- | --- | --- | --- | --- |
|  | Adult | Body | Child | Corridor | House | Limb | Number | Word |
| S1 | 0.751 | 0.701 | 0.746 | 0.715 | 0.670 | 0.652 | 0.708 | 0.774 |
| S2 | 0.810 | 0.749 | 0.837 | 0.755 | 0.651 | 0.690 | 0.640 | 0.777 |
| S3 | 0.786 | 0.763 | 0.822 | 0.743 | 0.608 | 0.719 | 0.747 | 0.811 |
| S4 | 0.809 | 0.745 | 0.794 | 0.747 | 0.566 | 0.682 | 0.535 | 0.727 |
| S5 | 0.779 | 0.691 | 0.787 | 0.740 | 0.605 | 0.687 | 0.633 | 0.799 |
| S6 | 0.779 | 0.721 | 0.812 | 0.759 | 0.632 | 0.654 | 0.601 | 0.771 |
| S7 | 0.757 | 0.714 | 0.788 | 0.726 | 0.574 | 0.643 | 0.647 | 0.789 |
| S8 | 0.799 | 0.804 | 0.840 | 0.802 | 0.646 | 0.699 | 0.616 | 0.797 |
| Mean | 0.784 | 0.736 | 0.803 | 0.748 | 0.619 | 0.678 | 0.641 | 0.781 |

**Table I.3:**
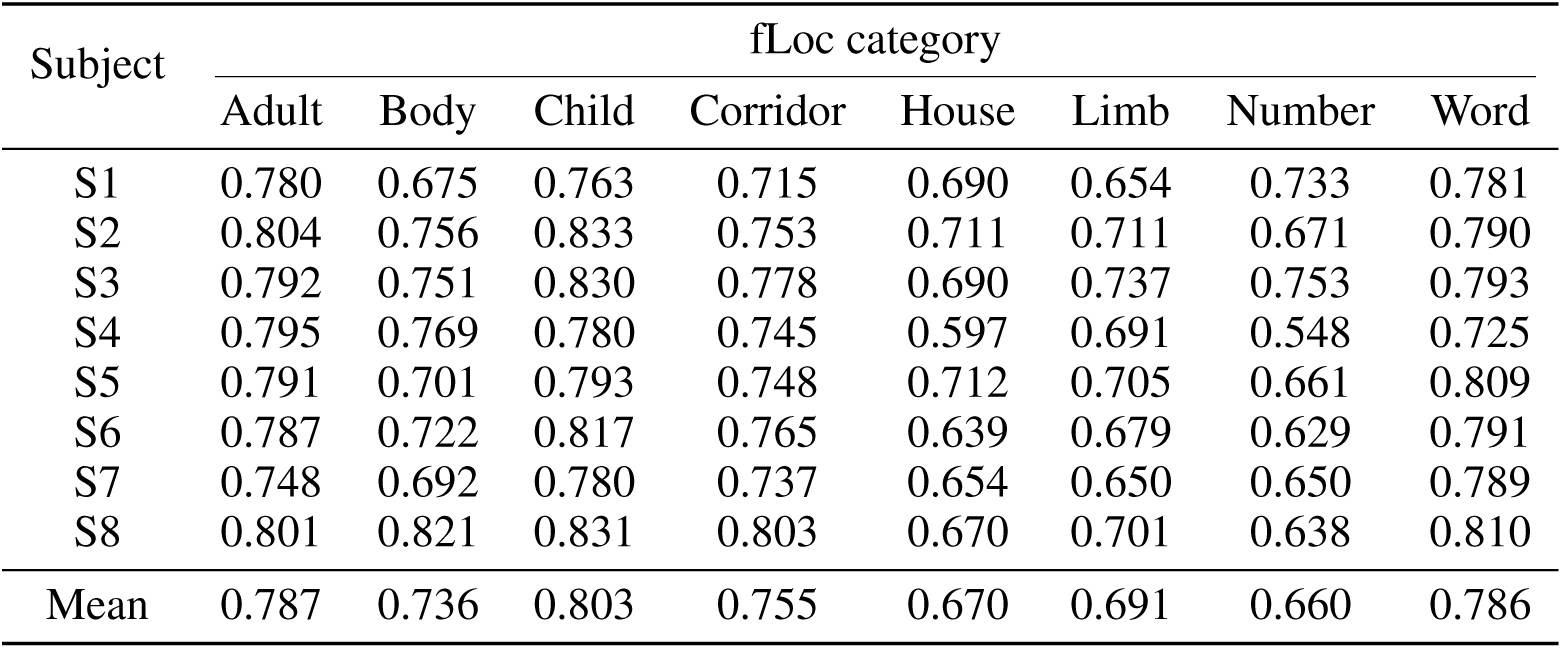
Cosine similarity between SAIL predicted and measured fLoc category responses for DeepSeek-VL-7B-Chat.

| Subject | fLoc category |  |  |  |  |  |  |  |
| --- | --- | --- | --- | --- | --- | --- | --- | --- |
|  | Adult | Body | Child | Corridor | House | Limb | Number | Word |
| S1 | 0.780 | 0.675 | 0.763 | 0.715 | 0.690 | 0.654 | 0.733 | 0.781 |
| S2 | 0.804 | 0.756 | 0.833 | 0.753 | 0.711 | 0.711 | 0.671 | 0.790 |
| S3 | 0.792 | 0.751 | 0.830 | 0.778 | 0.690 | 0.737 | 0.753 | 0.793 |
| S4 | 0.795 | 0.769 | 0.780 | 0.745 | 0.597 | 0.691 | 0.548 | 0.725 |
| S5 | 0.791 | 0.701 | 0.793 | 0.748 | 0.712 | 0.705 | 0.661 | 0.809 |
| S6 | 0.787 | 0.722 | 0.817 | 0.765 | 0.639 | 0.679 | 0.629 | 0.791 |
| S7 | 0.748 | 0.692 | 0.780 | 0.737 | 0.654 | 0.650 | 0.650 | 0.789 |
| S8 | 0.801 | 0.821 | 0.831 | 0.803 | 0.670 | 0.701 | 0.638 | 0.810 |
| Mean | 0.787 | 0.736 | 0.803 | 0.755 | 0.670 | 0.691 | 0.660 | 0.786 |

## J Evaluating Preferred Images with an Independent Encoding Model

We evaluate preferred images using a voxelwise encoding model trained on the Natural Scenes Dataset (NSD). Specifically, features extracted from a CLIP ViT backbone are fitted to individual voxel responses, enabling prediction of target ROI activations. This provides a response-based complement to the semantic comparison of image sets. The evaluation concerns model predictions, not additional fMRI measurements.

### Response evaluation

The encoding model predicts voxel responses from CLIP ViT image features. Averaging predictions across voxels within each ROI gives one response score per image. All NSD images viewed by the subject define the natural-image reference distribution, against which we compare the preferred image distributions. Within each ROI, we normalize scores to [*−*1, 1] using the minimum and maximum ROI-averaged predictions over the complete NSD image set.

### Preferred image selection

RAW retrieves five high-activation images from each voxel’s best matching feature, while SAEs-BrainMap retrieves five preferred images from its matched SAE unit. For SAIL, we select the highest-activation image from each of five matched units to keep the image count consistent.

### Statistical evaluation

We compare mean predicted ROI responses between SAIL and each other method using a one-sided test with 10,000 permutations. Benjamini–Hochberg FDR correction covers the multiple comparisons across ROIs and methods for each model and subject analysis. Stars indicate corrected significance levels: *∗q <* 0.05 and *∗ ∗ q <* 0.01.

## Figure conventions

Boxplots compare SAIL, SAEs-BrainMap, RAW, and the complete NSD image set in FFA, EBA, RSC, VWFA, and FOOD. Boxes span the interquartile range, center lines mark medians, and whiskers reach the most extreme observations within 1.5 interquartile ranges.

### J.1 QWEN2.5-VL-7B-INSTRUCT

**Figure J.1:**
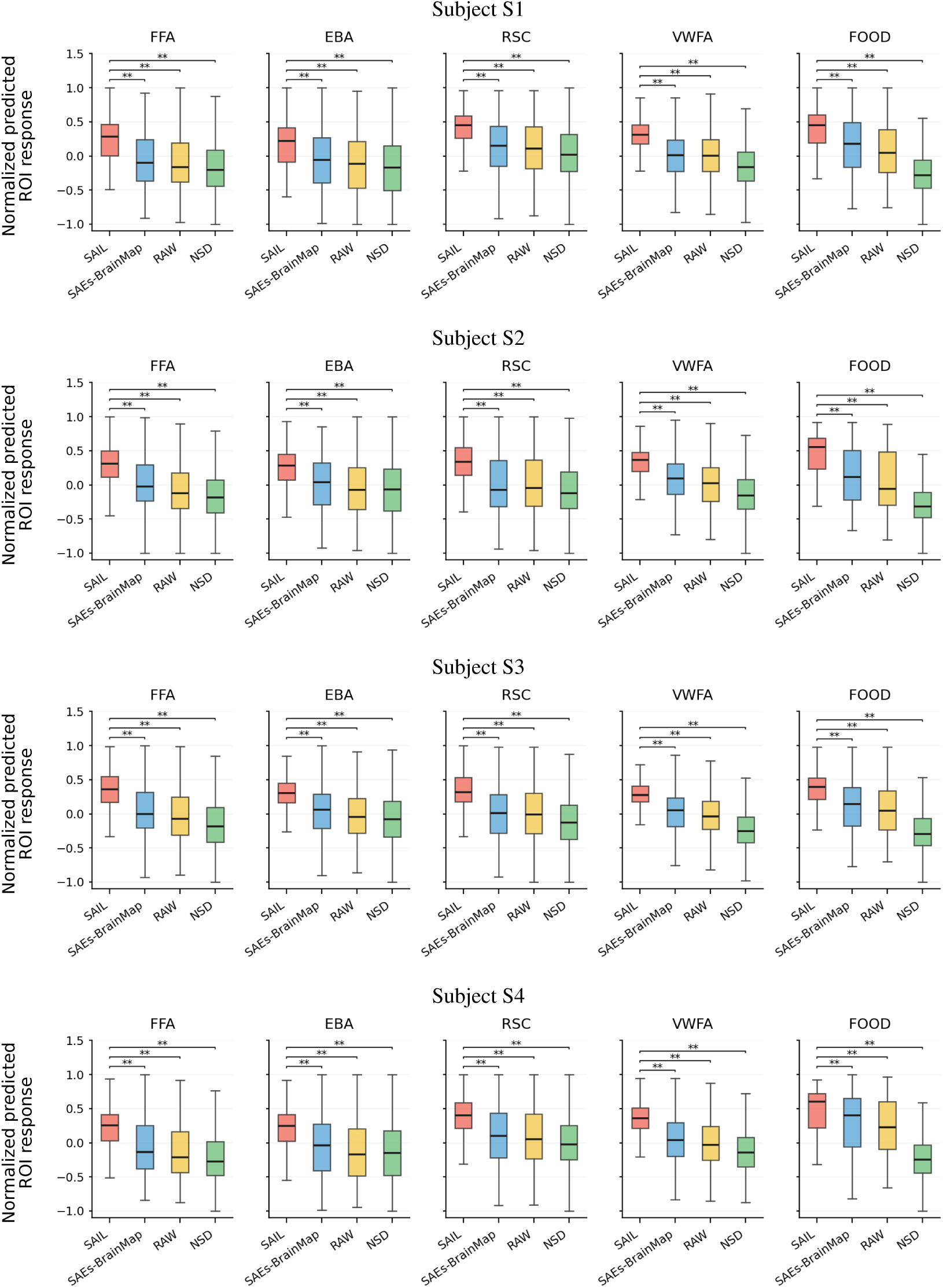
ROI response distributions predicted by the independent encoding model for Qwen2.5-VL-7B-Instruct, subject S1–subject S4. Stars indicate FDR-corrected significance from one-sided permutation tests: *∗q <* 0.05, *∗ ∗ q <* 0.01.

**Figure J.2:**
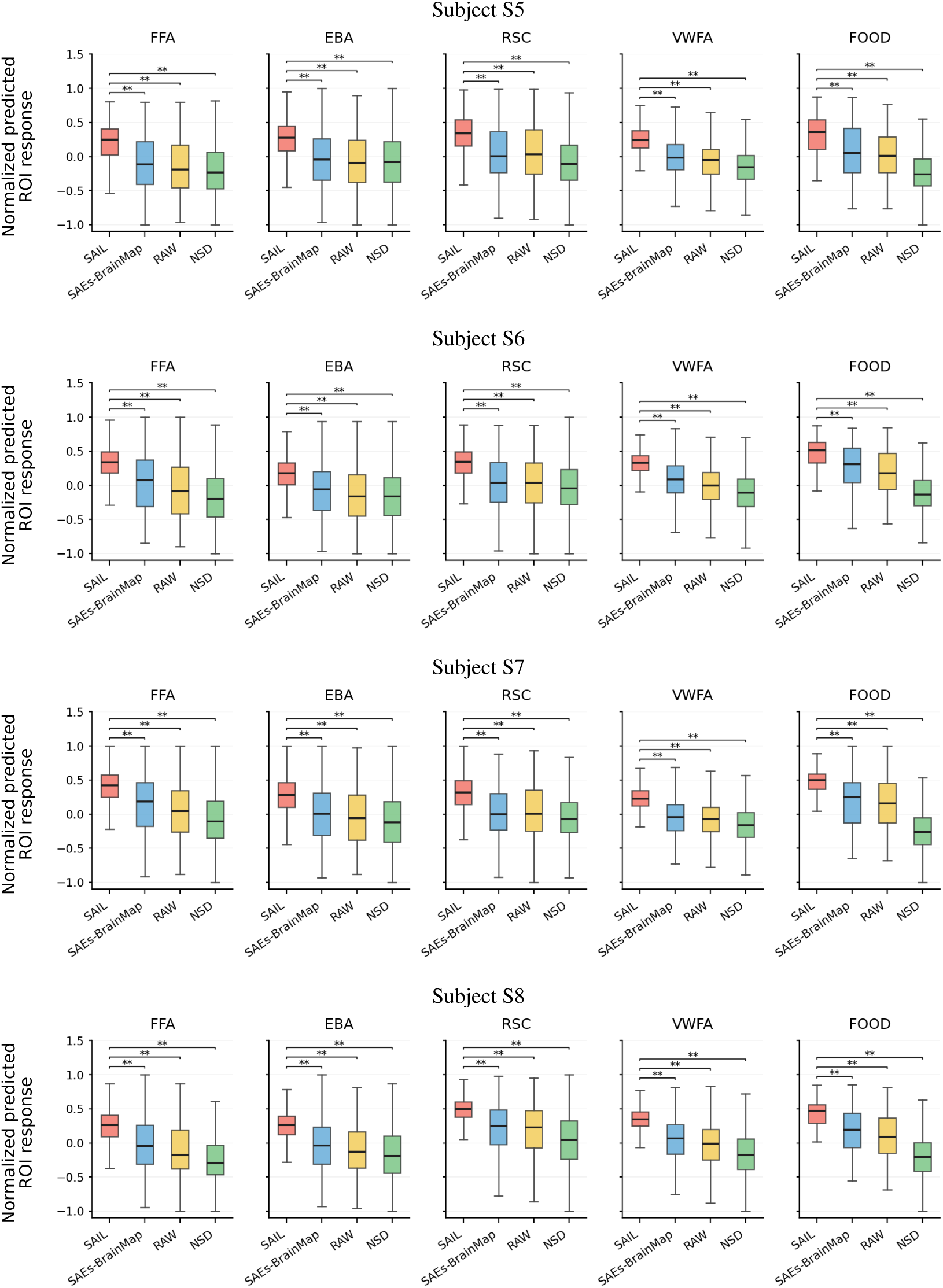
ROI response distributions predicted by the independent encoding model for Qwen2.5-VL-7B-Instruct, subject S5–subject S8. Stars indicate FDR-corrected significance from one-sided permutation tests: *∗q <* 0.05, *∗ ∗ q <* 0.01.

### J.2 LLAVA-V1.6-MISTRAL-7B

**Figure J.3:**
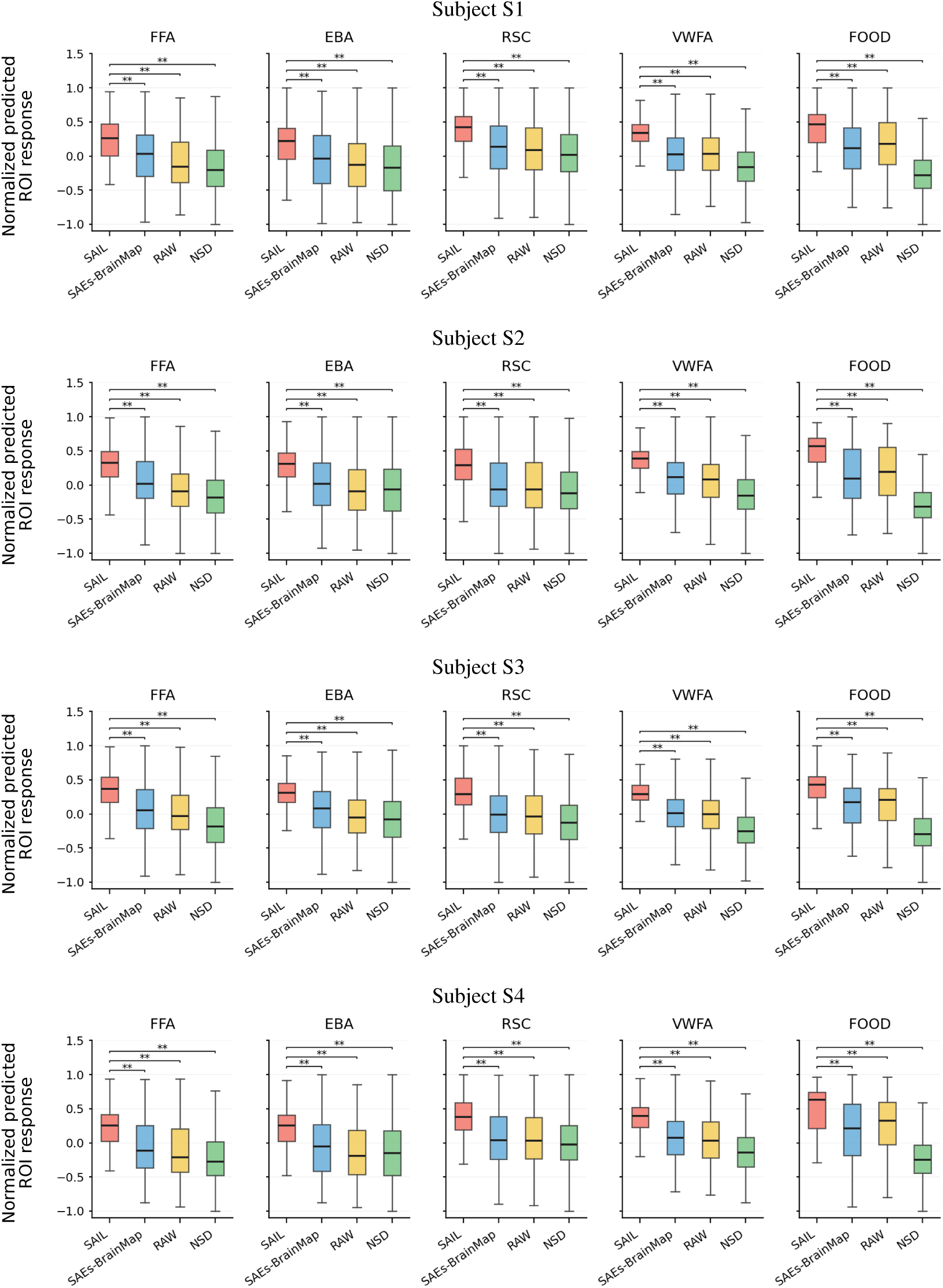
ROI response distributions predicted by the independent encoding model for LLaVA-v1.6-Mistral-7B, subject S1–subject S4. Stars indicate FDR-corrected significance from one-sided permutation tests: *∗q <* 0.05, *∗ ∗ q <* 0.01.

**Figure J.4:**
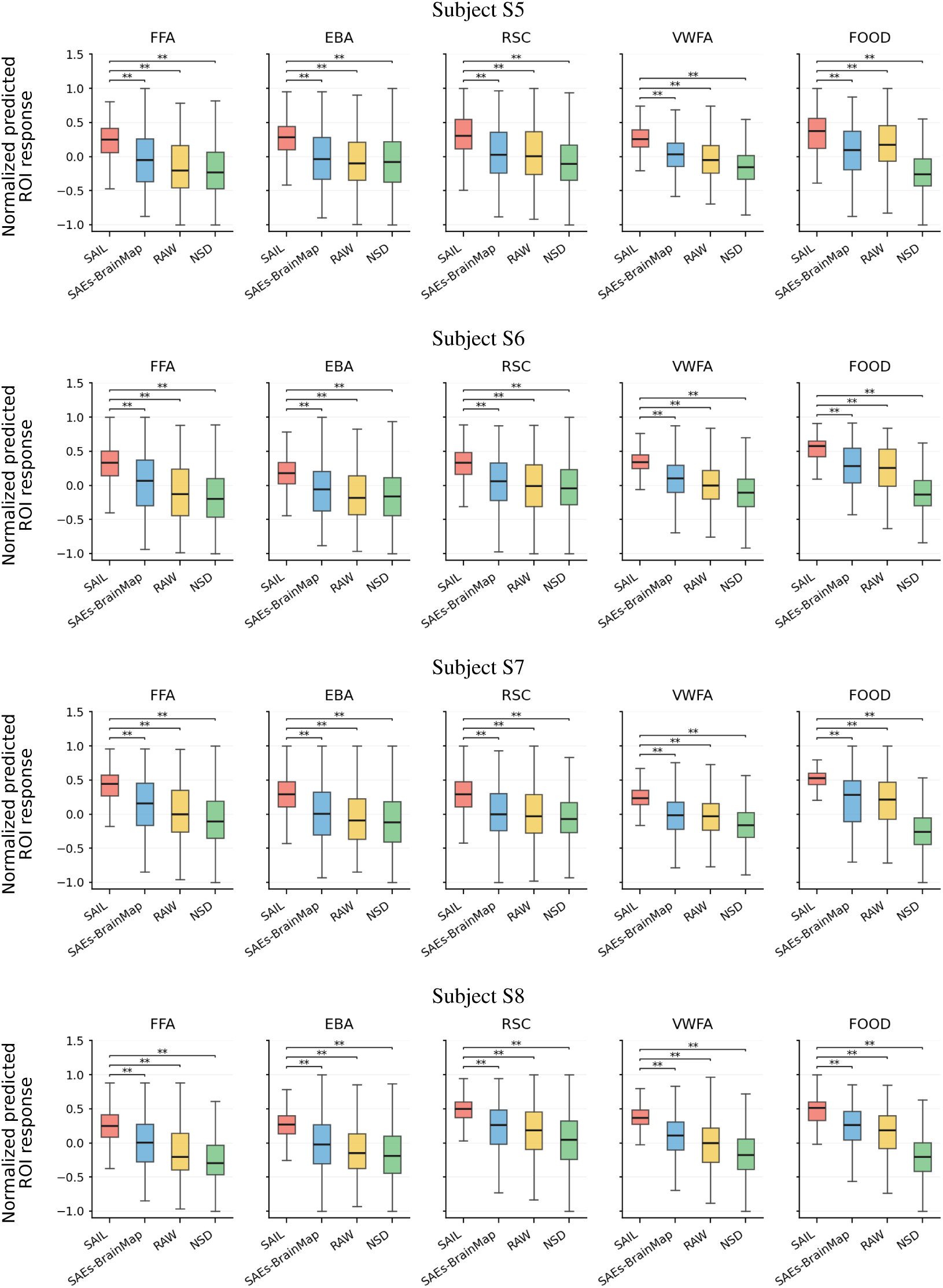
ROI response distributions predicted by the independent encoding model for LLaVA-v1.6-Mistral-7B, subject S5–subject S8. Stars indicate FDR-corrected significance from one-sided permutation tests: ∗q < 0.05, ∗ ∗ q < 0.01.

### J.3 LLAVA-V1.6-VICUNA-7B

**Figure J.5:**
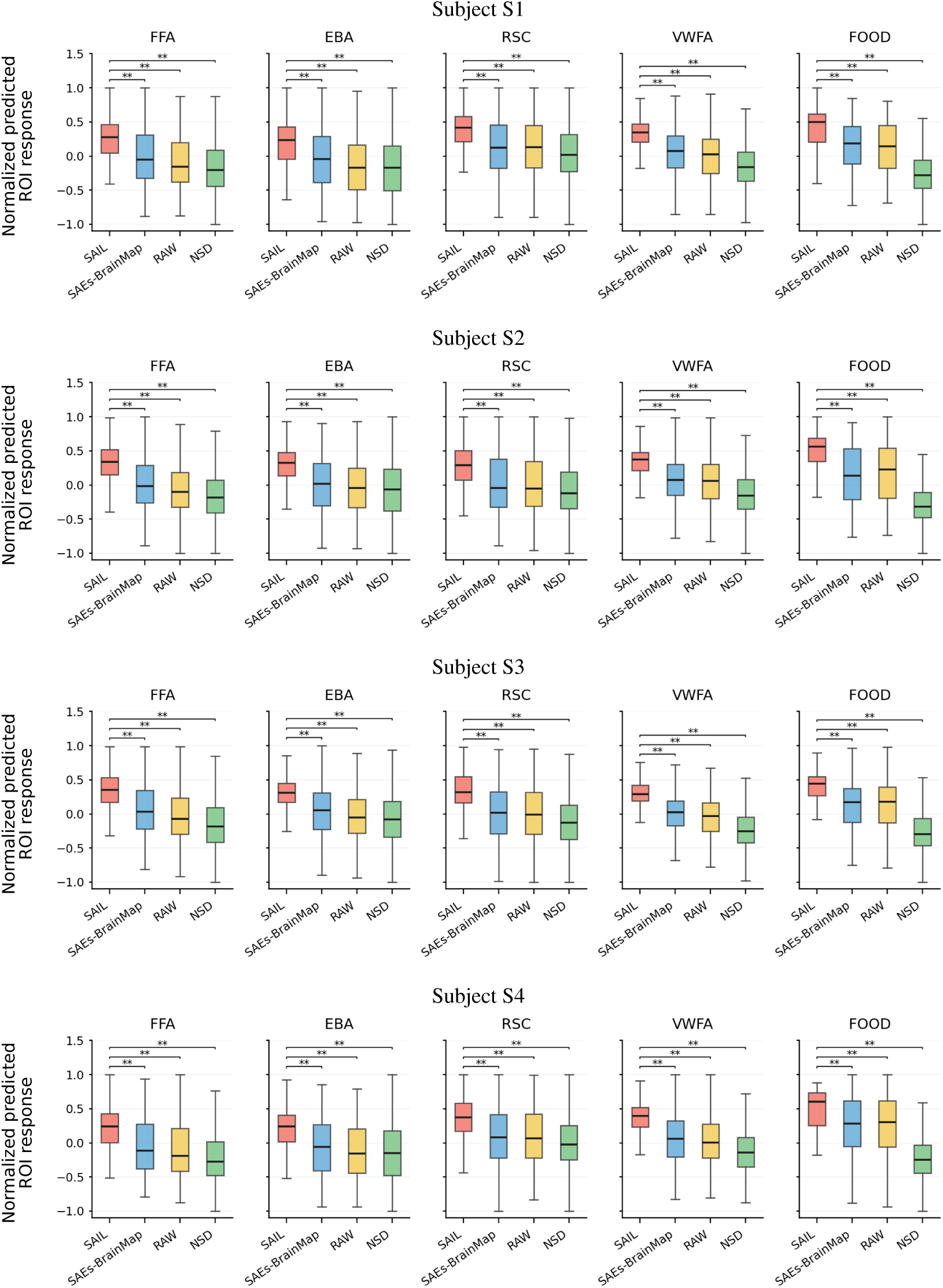
ROI response distributions predicted by the independent encoding model for LLaVA-v1.6-Vicuna-7B, subject S1–subject S4. Stars indicate FDR-corrected significance from one-sided permutation tests: *∗q <* 0.05, *∗ ∗ q <* 0.01.

**Figure J.6:**
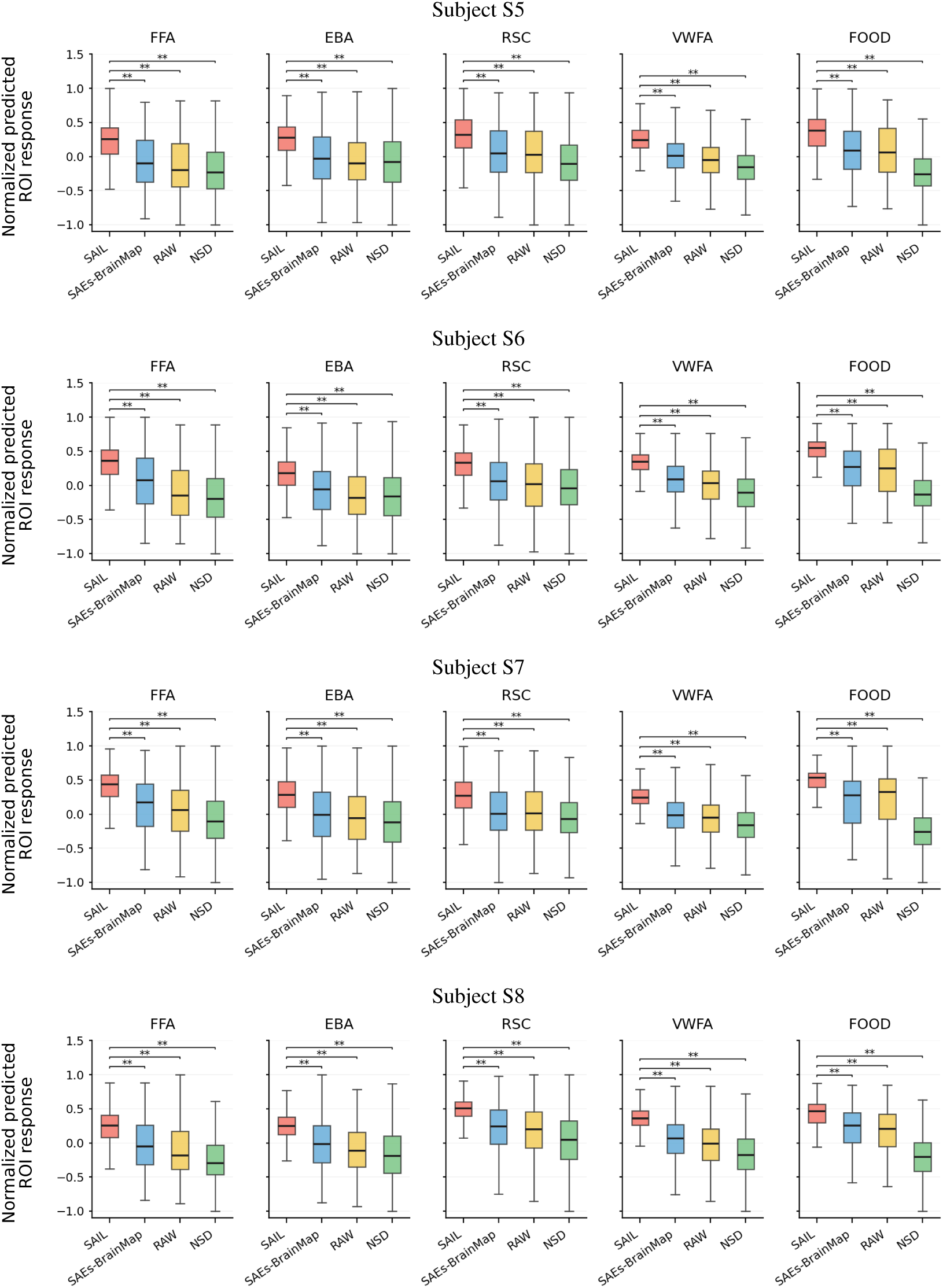
ROI response distributions predicted by the independent encoding model for LLaVA-v1.6-Vicuna-7B, subject S5–subject S8. Stars indicate FDR-corrected significance from one-sided permutation tests: ∗q < 0.05, ∗ ∗ q < 0.01.

### J.4 DEEPSEEK-VL-7B-CHAT

**Figure J.7:**
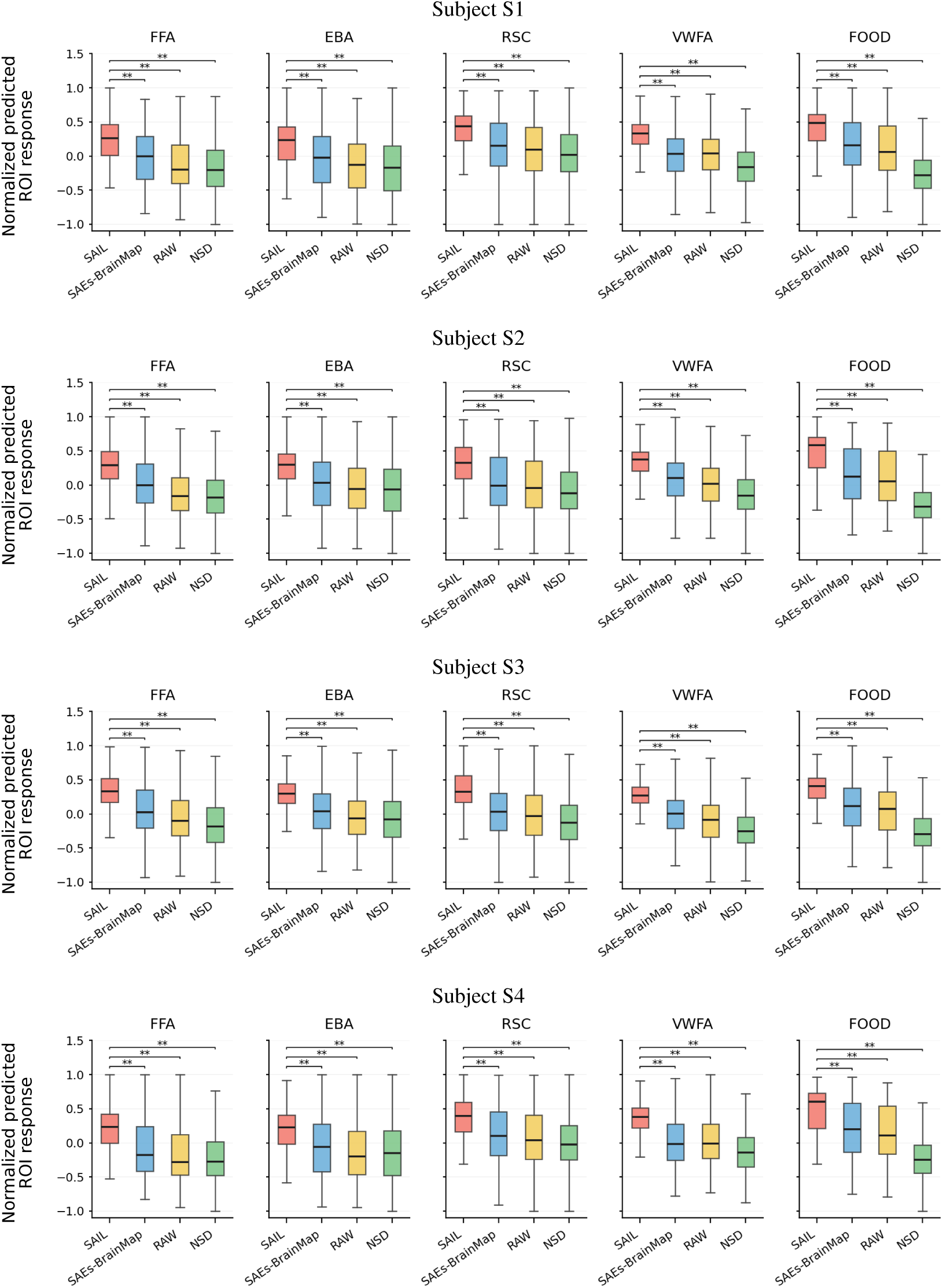
ROI response distributions predicted by the independent encoding model for DeepSeek-VL-7B-Chat, subject S1–subject S4. Stars indicate FDR-corrected significance from one-sided permutation tests: *∗q <* 0.05, *∗ ∗ q <* 0.01.

**Figure J.8:**
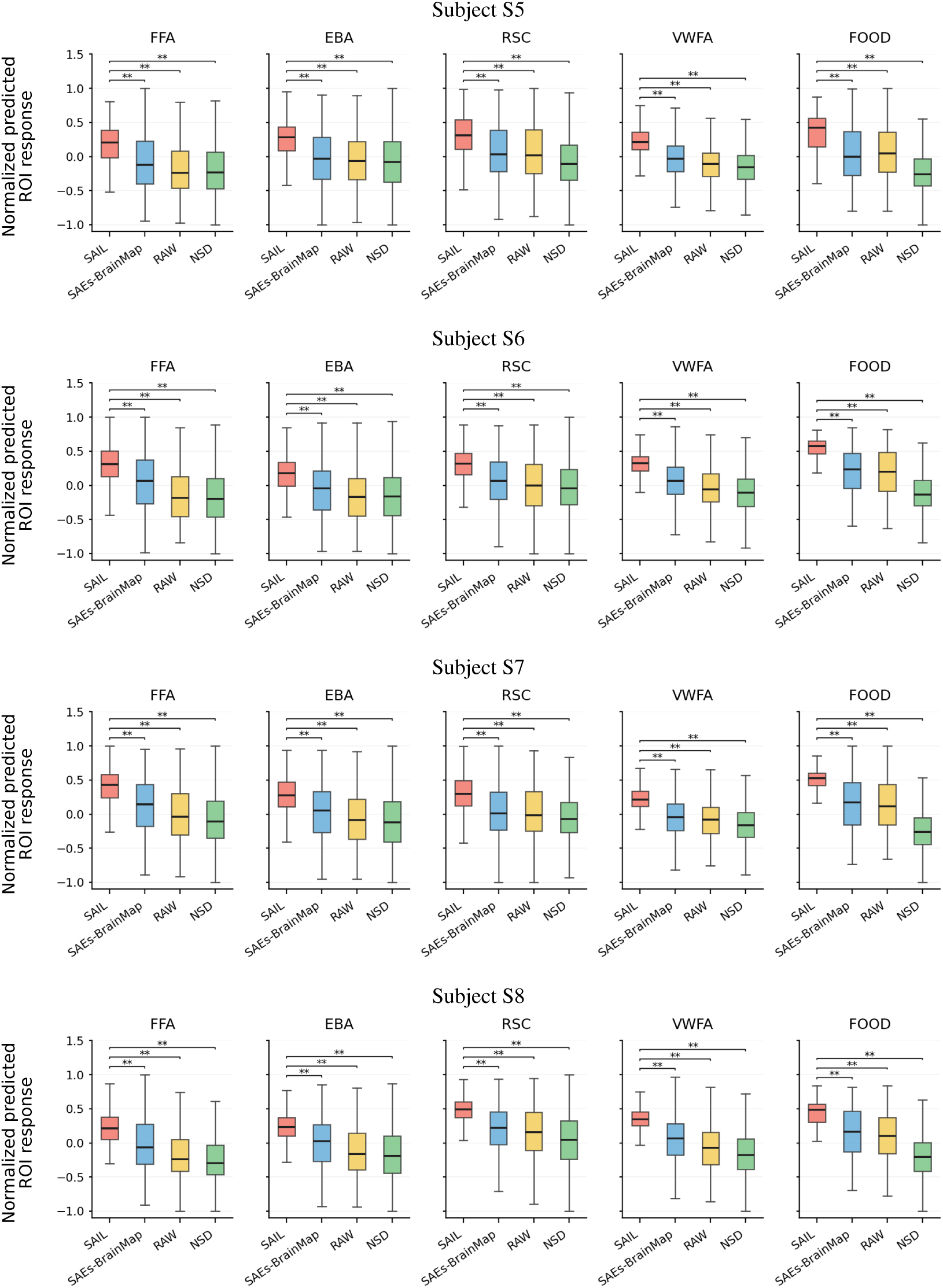
ROI response distributions predicted by the independent encoding model for DeepSeek-VL-7B-Chat, subject S5–subject S8. Stars indicate FDR-corrected significance from one-sided permutation tests: ∗q < 0.05, ∗ ∗ q < 0.01.

## K Construction of the Action-Related Vocabulary

### Vocabulary construction

We manually compiled a vocabulary to identify action-related semantic content in SAIL labels. The vocabulary was designed to capture both explicit actions and associated agents, equipment, and contextual cues in NSD/COCO images. Terms were organized into five operational groups: sports and activities, agents and groups, equipment and contexts, body movements and postures, and object interactions and everyday actions. We included selected singular and plural nouns and verb forms that could occur in SAIL labels. After deduplication, the vocabulary comprised 106 lowercase English terms. It was used as a fixed rule set during filtering, without language-model-based expansion or semantic-similarity matching.

### Label matching

Before matching, each SAIL label was stripped of leading and trailing whitespace and converted to lowercase. A label was considered a match only if the entire normalized string exactly matched one of the vocabulary terms. No substring matching, stemming, or lemmatization was applied. The five groups are used only to organize the vocabulary for presentation; group membership does not affect label matching or score computation. Because the vocabulary includes action-associated agents, equipment, and contexts in addition to explicit action words, a vocabulary match serves as an operational indicator of broadly defined action-related semantic content and does not, by itself, establish action selectivity.

### Complete vocabulary. Sports and activities (21)

baseball, basketball, competing, football, frisbee, game, golf, golfing, hockey, match, parade, performance, performing, play, playing, race, racing, soccer, softball, tennis, volleyball. **Agents and groups (22):** athlete, athletes, biker, bikers, crowd, dancer, dancers, golfer, group, motorcyclist, player, players, rider, riders, skier, skiers, skateboarder, snowboarder, snowboarders, surfer, surfers, team. **Equipment and contexts (22):** ball, bat, bicycle, bike, board, court, field, glove, helmet, horse, horses, kite, kites, motorcycle, racket, skate, skateboard, ski, skis, snowboard, stadium, surfboard. **Body movements and postures (23):** climb, climbing, dance, dancing, jump, jumping, ride, riding, run, running, sit, sitting, skating, skiing, snowboarding, stand, standing, surfing, swim, swimming, swinging, walk, walking. **Object interactions and everyday actions (18):** carrying, catch, catching, cooking, cutting, drinking, eating, feeding, grabbing, hitting, holding, kicking, pulling, pushing, reaching, throw, throwing, touching.

